# Probing Action Potential Generation and Timing under Multiplexed Basal Dendritic Computations Using Two-photon 3D Holographic Uncaging

**DOI:** 10.1101/2022.09.26.509562

**Authors:** Shulan Xiao, Saumitra Yadav, Krishna Jayant

## Abstract

Basal dendrites of layer 5 cortical pyramidal neurons exhibit Na^+^ and NMDAR spikes, and are uniquely poised to influence somatic output. Nevertheless, due to technical limitations, how multibranch basal dendritic integration shapes action-potential output remains poorly mapped. Here, we combine 3D two-photon holographic transmitter-uncaging, whole-cell dynamic-clamp, and biophysical modeling, to reveal how synchronously activated synapses (distributed and clustered) across multiple basal dendritic branches impacts action-potential generation – under quiescent and *in vivo* like conditions. While dendritic Na^+^ spikes promote milli-second precision, distributed inputs and NMDAR spikes modulate firing rates via axo-somatic persistent sodium channel amplification. Action-potential precision, noise-enhanced responsiveness, and improved temporal resolution, were observed under high conductance states, revealing multiplexed dendritic control of somatic output amidst noisy membrane-voltage fluctuations and backpropagating spikes. Our results unveil a critical multibranch integration framework in which a delicate interplay between distributed synapses, clustered synapses, and axo-somatic subthreshold conductance’s, dictates somatic spike precision and gain.

## INTRODUCTION

Cortical pyramidal neurons in layer 5 (L5) respond with millisecond precision to sensory inputs (London et al., 2010; Tiesinga et al., 2008). Here, rapidly time-varying feedforward streams, predominantly synapsing onto L5 basal dendrites, cause somatic action potentials (AP) with millisecond resolution precision. From a functional perspective, precise control of spike timing as well as burst rate is critical for spike-timing-dependent plasticity (Duguid and Sjöström, 2006; Kampa et al., 2007), ensuring activation of apical calcium (Larkum et al., 1999; Pérez-Garci et al., 2006) and basal dendritic NMDAR (N-methyl-D-aspartate receptor-dependent) spikes (Kampa and Stuart, 2006), and enabling calcium-activated burst mode signaling - a hallmark of optimal cortical coding (Takahashi et al., 2020). However, the stochastic nature of synaptic transmission, dendritic filtering, and high-levels of background noise imply that signal integration and eventual AP generation will necessarily be a variable process. Here, background noise consists of constantly active conductances and sharp voltage fluctuations. Computational studies (Destexhe et al., 2003; Hô and Destexhe, 2000) backed by experiments show that under high-conductance conditions, the presence of sharp voltage fluctuations can in fact, reduce spike jitter and improve AP precision (Ariav et al., 2003; Fernandez et al., 2011; Mainen and Sejnowski, 1995; Shu et al., 2003). Dendrites, with their unique morphology, varied resistance, and myriad of nonlinear regenerative transformations, are believed to be a locus of such sharp fluctuations and play a key role in determining accurate AP generation. This mechanism is well complemented by recent *in vivo* imaging efforts in which widespread multibranch dendritic activity results in highly coupled somato-dendritic dynamics, and the highly-fluctuating conditions they generate could play a critical role in AP precision and burst generation (Beaulieu-Laroche et al., 2019; Connelly and Stuart, 2019; Destexhe et al., 2003; Kerlin et al., 2019). Studies have also alluded to the role of intrinsic conductance’s in controlling AP gain (Hsu et al., 2018; Yue et al., 2005), yet, the interplay with synaptic input patterns, widespread dendritic regenerative events, and noisy conductance’s remains poorly mapped. This interplay is critical to our understanding of neural selectivity and plasticity (Legenstein and Maass, 2011), but has remained experimentally intractable so far (Ujfalussy et al., 2018).

Regenerative dendritic events, mainly NMDAR spikes and Na^+^ spikes, generated by synchronously activated synaptic clusters on thin basal and tuft dendrites (Antic et al., 2010; Larkum and Nevian, 2008; Losonczy et al., 2008) influence sensory coding (Gambino et al., 2014; Kumar et al., 2018; Lavzin et al., 2012; Palmer et al., 2014), AP generation (Ariav et al., 2003; Losonczy et al., 2008; Palmer et al., 2014), and plasticity (Lee et al., 2016; Weber et al., 2016). These synaptic clusters span 8 µm to 15 µm stretches and can be spread across multiple dendrites and interspersed alongside distributed inputs (Gökçe et al., 2016; Hill et al., 2013; Jia et al., 2010; Markram, 1997; Scholl et al., 2017; Takahashi et al., 2012). It is now becoming clear that such clusters of coactive synapses with unique subcellular distributions, possibly originating from the same presynaptic assembly, are a major factor in determining the nature of regeneration and sensitivity of a single pyramidal neuron (Gökçe et al., 2016; Iacaruso et al., 2017; Ju et al., 2020; Rah et al., 2013; Scholl et al., 2017). The input-output relationship across these thin basal dendrites follows the well accepted two-layer-model of computation (Jadi et al., 2014; Poirazi et al., 2003) with a stark location-dependence (Behabadi et al., 2012). Here, synchronously activated inputs result in a sigmoidal input-output transformation when clustered, a linear to sub-linear relationship when distributed (Branco and Häusser, 2011; Major et al., 2008), exhibit a NMDAR dependence (Branco and Häusser, 2011; Hill et al., 2013; Lafourcade et al., 2022; Major et al., 2008; Schiller et al., 2000), and play a crucial role in axo-somatic integration (Antic et al., 2010; Major et al., 2013). .

Basal dendrites of L5 pyramidal neurons, which receive a diverse range of clustered feed-forward (Petreanu et al., 2009) and intra-cortical synaptic inputs (Gökçe et al., 2016; Markram, 1997; Sjöström and Häusser, 2006) are thus uniquely poised to exert a potent influence on somatic action potential dynamics (Dembrow and Spain, 2022; Markram, 1997; Williams, 2004). With this backdrop it is interesting to note, albeit in a different class of neurons, that previous studies by Haag and Borst (Haag and Borst, 1996), Schiller and colleagues (Ariav et al., 2003) and subsequently by Losonczy et al., (Losonczy et al., 2008), converged onto the same underlying principle that dendritic Na^+^ spikes are in fact an important determinant of spike precision at the soma. However, these studies did not differentiate between the contributions of clustered and distributed synchronous synaptic inputs. Furthermore, the mechanisms of precision and burst generation under multibranch conditions were not probed, and experiments were performed under quiescent low-noise conditions. What is the impact of cluster-evoked dendritic nonlinearities to AP timing and rate under noisy backgrounds and increased conductance? Is there a critical number of synaptic inputs that must be co-active to impact AP output under *in vivo* like conditions? Do backpropagating spikes reduce the efficacy of synaptic cluster-evoked somatic control? Unravelling how coincident synaptic inputs (clusters and distributed inputs) across multiple basal dendrites integrate under conditions of quiescence and increased background conductance will allow us to address these outstanding questions.

Previous efforts to unravel the role of multibranch integration in pyramidal neurons have been scarce. Reports focusing on assaying the role of multisite/multibranch dendritic inputs neuronal integration have used multisite focal theta stimulation (Kumar et al., 2018), acousto-optic-deflectors (AODs) based uncaging (Losavio et al., 2009), and single (Lutz et al., 2008; Yang et al., 2014) or two-photon holographic uncaging (Go et al., 2019; Nikolenko et al., 2008) with phase-only spatial-light-modulators. While multisite theta stimulations undoubtedly assist in generating dendritic spikes, the ability to precisely stimulate clusters and distributed inputs in synchrony and across space is not possible. AODs on the other hand, offer higher positioning speeds but suffer from spatio-temporal distortion of ultrafast laser pulses, and exhibit a wavelength-dependent low diffraction efficiency leading to small field of views, precluding 3D uncaging to date (Losavio et al., 2009). Two-photon uncaging of transmitters using a set of galvanometers allows for precise stimulation of individual and synaptic clusters respectively (Branco and Häusser, 2011; Harnett et al., 2012; Lafourcade et al., 2022; Losonczy et al., 2008; Weber et al., 2016). Galvanometric mirrors can be programmed to allow for fast beam steering and rapid repositioning (∼ 100 *μ*s) with little to no power loss, however, they are limited to a single focal plane and cannot assay three-dimensional synaptic distributions (Gasparini et al., 2004). While single photon holographic uncaging suffers from poor point-spread-functions (PSFs) (Lutz et al., 2008), two-photon holographic uncaging promises to overcome these limitations, yet previous efforts have fallen short of experimental demonstrations of the full 3D capability (Go et al., 2019; Nikolenko et al., 2008).

Here, we combine a spatial-light-modulator (SLM) with tabletop pulse compression optics into a conventional two-photon microscope to enable 3D caged-transmitter uncaging in thick scattering tissue. We combine this holographic uncaging approach with whole-cell electrophysiology, dynamic clamp, and computational modeling to elucidate the mechanisms underlying precise AP timing and burst control under synchronous synaptic streams spread across multiple basal dendritic branches with a depth range ∼40 µm (max 70 µm). Using dynamic clamp, we examine the factors underlying AP generation both under quiescent and noisy *in vivo* like states, and provide a quantitative biophysical model of how spike timing and bursting is generated as a function of dendritic nonlinearities and distributed synaptic inputs. Specifically, we reveal a critical interaction between dendritic spikes and persistent sodium currents at the axo-somatic junction that promotes bursting, while stochastic channel activity promotes precise spike timing. By examining the role of high-conductance states on multibranch integration, we reveal that precision and rate control are maintained under conditions of high conductance, noisy membrane-potential fluctuations, and are not affected by backpropagating action potentials.

## RESULTS

### 3D Two-photon Holographic Uncaging: Optical and Physiological Characterization

Our overall system comprises a commercial two-photon microscope integrated with an SLM module, custom-designed table-top compression optics, a Michelson interferometer, and dual laser lines for simultaneous imaging and uncaging (**Figure 1A (i)**). The set-up is capable of 3D stimulation (**Figure 1A (i), inset, top**) with power weighting for each point independently (**Figure 1A (i), inset, bottom, see Methods**). The point spread function of our SLM-based system (0.6 µm lateral, 2.5 µm axial, **Figure 1A (ii)**) measured using 100nm fluorescent beads was found to be identical to that with just galvanometric mirrors (**Figure S1A**). While the lateral PSF is in line with previous experimental measures (Ellis-Davies, 2019), the axial PSF was slightly higher than previously reported values (∼ 1.3 µm – 2 µm axial) (Harvey and Svoboda, 2007; Matsuzaki et al., 2001; Smith et al., 2003). We attribute our slightly higher axial PSF to the marginally underfilled objective (0.8 NA) in our design, which is critical to maximize power transmission by the SLM. This is done to obtain an optimal tradeoff between power per spine and a desired resolution. The combined use of the SLM and galvanometric mirrors allowed us to reposition the holographic FOVs rapidly (**Figure S1B**) while ensuring diffraction-limited synaptic stimulation in 3D with a desired number of beamlets. Secondly, the use of a modular custom-designed prism-based compressor allowed us to achieve sub-100 fs pulse widths at the sample plane for efficient uncaging (**Figure 1A (iii)**), albeit slightly higher than that with galvanometric mirrors (**Figure S1C**).

**Figure 1:**
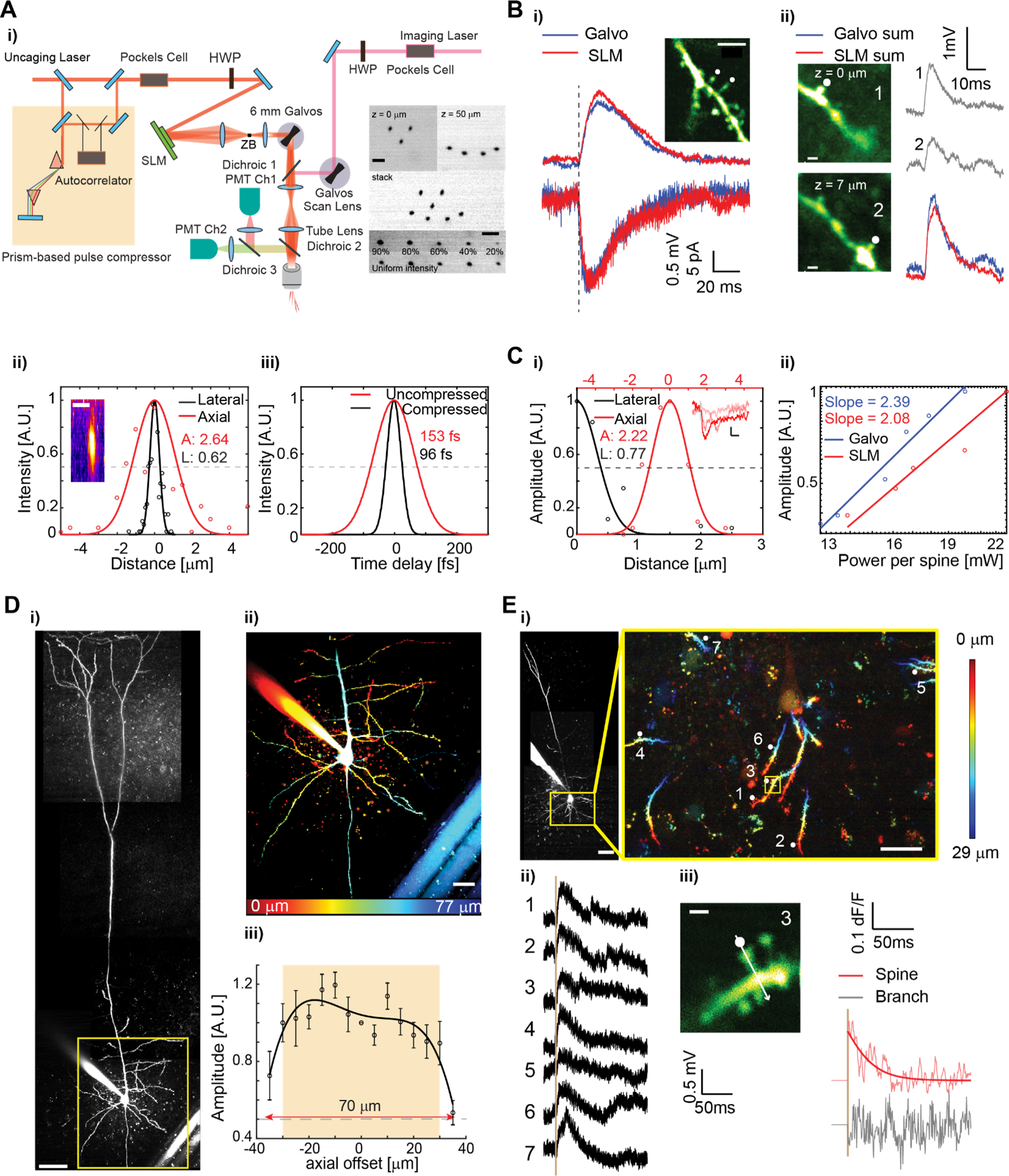
3D Two-photon spatial light modulator (SLM) based uncaging: **(A, i)** Schematic depicting the layout of the two-photon microscope with imaging and uncaging paths including the SLM module. (Right inset) Examples of holographic patterns (top) and holographic spots with independent power weighting (bottom). Scale bars, 5 µm. **(A, ii)** Point spread function (PSF) of a first order SLM beamlet measures with fluorescent beads. **(A, iii)** Pulse width measures with the SLM module in the uncaging path. (Inset) x-z projection of a 100nm fluorescent bead imaged with a first order SLM beam. Scale bars, 2 µm. **(B)** Comparison of holographic uncaging vs. galvanometric uncaging responsivity. **i)** EPSP and EPSC’s induced by uncaging across 2 spines (the time of activation marked with the dash line) with galvanometric scanners (blue) and SLM (red). The inter-stimulus-interval for galvanometer-based stimulation is 0.3 ms. Scale bars, 5µm. **ii)** An example depicting the response upon simultaneous uncaging of 2 spines with the SLM located across different depths (red), compared to the linear sum of the individual responses from each spine under galvanometric uncaging (blue). Galvanometric uncaging response for each spine is shown in gray. Scale bars, 1µm. **(C, i)** physiological PSF under holographic uncaging measured as the amplitude of unitary EPSP and unitary EPSC’s as a function of the lateral (black) and axial (red) distance between the uncaging location and the spine head, fitted with Gaussian curves. Inset: Unitary EPSC transients as a function of shifting holographic spot offsets from the spine head (0, 0.25, 0.5 µm) scale bars, 5ms (horizontal) and 5pA (vertical). **(C, ii)** Log-log plot of unitary EPSP amplitudes plotted as a function of incident laser power for galvanometric (blue) and SLM (red) uncaging. **(D)** Uncaging efficiency across depth. **i)** An example L5 pyramidal neuron. Scale bars, 50 µm. **ii)** Magnified view of basal dendrites from region shown in (A). Basal arbors are distributed across ∼ 70 µm of depth. Scale bars, 20µm. **iii)**The unitary EPSP amplitude (normalized to amplitude when axial offset is 0) as the function of the axial offset of the 1^st^ order SLM beam, N = 3 cells. A near uniform uncaging response is observed for two beamlets separated axially by ∼70 µm (FWHM). **(E, i)** An example L5 pyramidal neuron and magnified view of the basal dendrites. Scale bars, 50µm (main) and 20µm (inset). **(E, ii)** Scan-less single spine unitary EPSP responses from 7 individual basal dendritic spines. **(E, iii)** A Ca^2+^ line scan performed across spine head and adjacent dendritic shaft of spine number 3. Scale bars, 1µm.

Next, we measured the physiological PSF of SLM-based uncaging and compared it with galvanometric uncaging. We found that glutamate uncaging responses (1.5 mM DNI-glu; 1 ms pulses; **see Methods for details**) with both the SLM and galvanometric mirrors on two spines located in the same plane and two different planes separated by 7 µm’s were identical (**Figure 1B**). Here, galvanometric uncaging was performed at both the focal planes independently and compared to the holographic uncaging response. The physiological PSF of the SLM (**Figure 1C (i)**) was found to be indistinguishable from the measures using beads (**Figure 1A (ii)**), and resulted in the same excitatory-post-synaptic-potential (EPSP) and excitatory-post-synaptic-current (EPSC) kinetics and amplitudes in comparison to galvanometric scanning (**Figures S1D, S1E**). The power needed to evoke a unitary response (∼0.5 mV) from a single spine with the SLM engaged was 15mW – 17mW with pulse compression (**Figure 1C, (ii)**) and this was only slightly higher than the power needed for uncaging with galvanometric mirrors. Importantly, pulse compression did not change the physiological PSF measures (**Figure S1F**). This ensured stimulation of spines with no observable photodamage even after repeated stimulations (**Figure S1G**; **also see Methods**).

L5 pyramidal neuron basal dendrites project three dimensionally and are distributed across a depth of approximately 50 µm (**Figure 1D (i, ii)**). Since the diffraction efficiency of the SLM decreases with increasing angle of beam steering, we calibrated the axial limits of the SLM before we encountered an appreciable power roll-off. To this end, we centered a single holographic spot (beamlet) on a spine head of interest and uncaged while gradually defocusing the zeroth order focal-plane of the objective axially. The amplitude of spine evoked EPSPs measured at the soma remained identical across an axial range of ± 35um (**Figure 1D (iii)**) resulting in a uniform uncaging FOV across a 70 µm depth, which largely covers the L5 PN basal dendritic arborization in acute brain slices. We then performed scan-less holographic uncaging across several spines spread across diametrically opposite basal arbors encompassing a 100 µm * 100 µm * 35 µm FOV and found that each synapse was uncaged with similar efficiency and specificity (**Figure 1E**). Critically, simultaneous calcium imaging across a chosen spine reflected compartmentalized calcium signaling restricted to the spine head and not the dendritic shaft, suggesting minimal to no off-target activation. However, when clusters of synapses were activated, calcium signals in dendrites reflected the cooperativity across clustered inputs (**Figure S1H**).

### Holographic uncaging evoked dendritic nonlinearities dictate somatic spike timing and gain

Basal dendrites of L5 pyramidal neurons exhibit a rich repertoire of local nonlinearities including 2D sigmoidal input-output transforms. They also have a high input impedance and a large space constant (Nevian et al., 2007). A key component underlying these nonlinearities are clusters of co-active closely spaced synapses which via NMDAR cooperativity lead to local dendritic spikes (d-spikes). Using 3D two-photon holographic uncaging of synaptic clusters across multiple basal dendrites of a single L5 pyramidal neuron, we observed distinct nonlinear input-output transformations as a function of number of synapses stimulated simultaneously (**Figure 2A**), characteristic of a sigmoidal input-output pattern (**Figure S2A and S2B**). Here, cells were hyperpolarized from rest to reveal the full length of the nonlinear d-spike without evoking a somatic AP. It is important to note that given the maximum power available at the sample plane (∼180 mW), we were able to stimulate a maximum of 12 spines in a cluster with high-efficiency, and up to 36 spines spread across 3 groups along three separate dendritic arbors with the only delay being the 3 ms needed between phase masks. While some basal dendritic clusters exhibited a fast-rising Na^+^ spike characterized by a high dV/dt followed by a large NMDAR dependent plateau (**Figure 2A, red and magenta**) (strong cluster), several clusters revealed a purely NMDAR dependent signal (**Figure 2A, black)** (weak cluster). The NMDAR plateau potential was found to be dependent on AP5, while the Na^+^ spike was critically dependent on Tetrodotoxin (TTX) (**see Figure S2 C-E**). We then slowly ramped the resting membrane potential (RMP) of the cell via the patch electrode and observed that when the RMP was close to threshold, dendritic nonlinearities exhibiting the fast Na^+^ spike resulted in a precise AP output with millisecond precision and very low jitter (**Figure 2B, top**). However, d-spikes reflecting pure NMDAR events led to AP output with much larger variability with high jitter (**Figure 2B, middle**). Importantly, a set of Na^+^ d-spikes followed by large NMDAR plateaus also resulted in precise bursts of APs with very low jitter (**Figure 2B, bottom**). Na^+^ spikes characterized by the fast dV/dt component showed no observable trend with respect to overall d-spike amplitude, but rather a spread (**Figure 2C**). These strong d-spikes also exhibited a weak location dependence along the basal dendrites with Na^+^ spikes, occurring between 50 to 150um away from the soma (**Figure 2D**). But they were critically dependent on the resting membrane potential (**Figure 2E**). In comparison, NMDAR events alone exhibited a large jitter, i.e. standard deviation of spike timing (**Figure 2F, column i**) (Mann–Whitney–Wilcoxon test, N = 11 strong branches and 16 weak branches from 20 cells, p < 0.001), imprecise onset (**Figure 2F, column ii**) (Mann–Whitney–Wilcoxon test, N = 14 strong branches and 28 weak branches from 20 cells, p < 0.001). Taken together, the results shown in **Figures 2A-2F** would suggest that while spike timing is critically dependent on Na^+^ d-spikes, NMDAR spikes lead to variable jitter, and both influence AP output in an RMP-dependent fashion.

**Figure 2:**
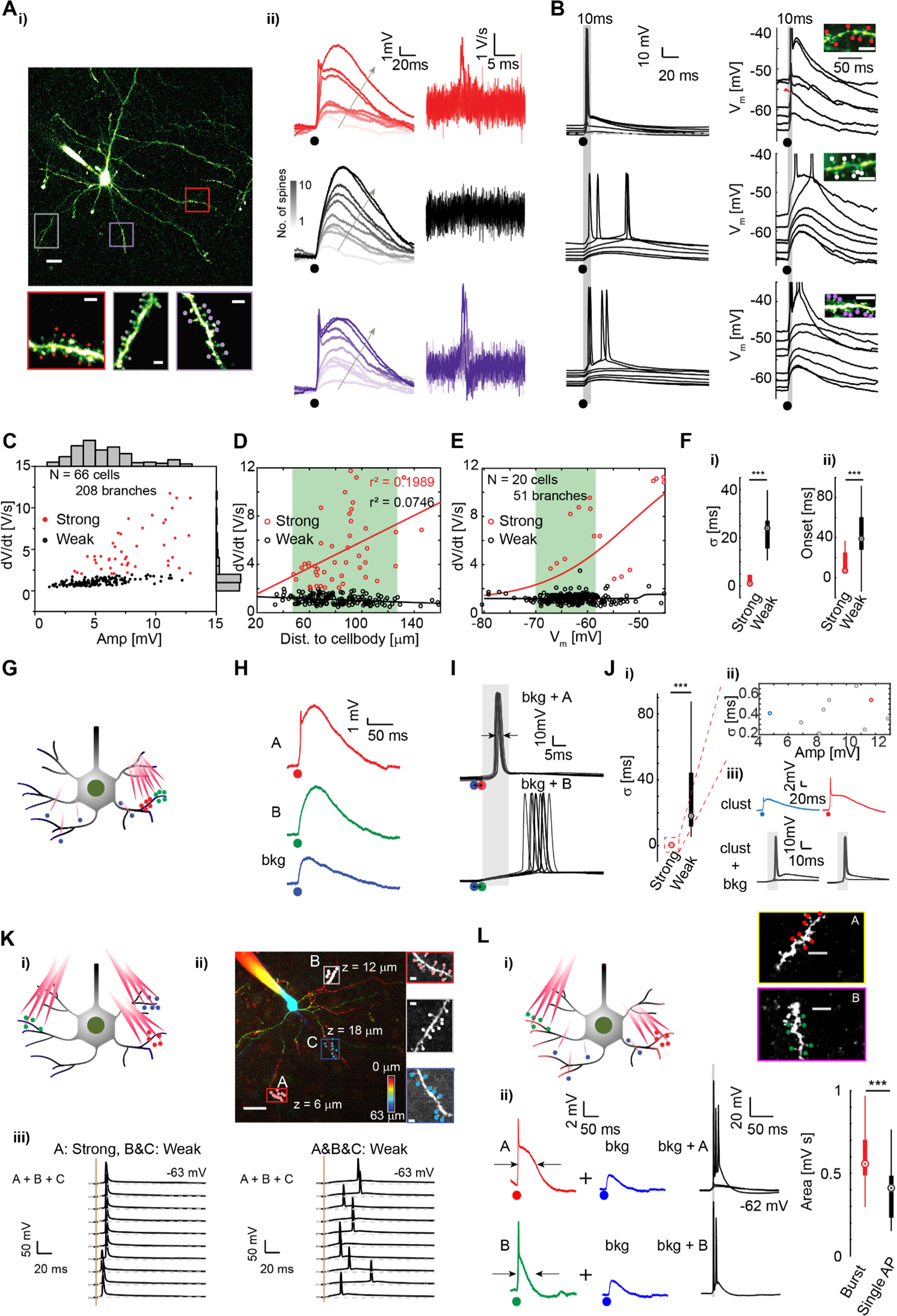
Clustered synaptic activation-evoked action potential dynamics in a multi-branch framework. **(A, i)** An L5 pyramidal neuron (scale bars, 20 µm) and three basal dendritic segments (scale bars, 2 µm) across which clustered synapses were stimulated. **(A, ii)** Holographic uncaging-evoked potentials measured at the soma as a function of increasing number of spines within each branch. **(A, ii, right)** dV/dt corresponding to uncaging waveforms. **(B)** Holographic uncaging response of synaptic clusters as a function of varying RMP elicited via current-injection through the whole-cell patch electrode (uncaging shown by black dot). **(B, right)** magnified view of the somatic responses during cluster activation (6-8 spines, inset). (Inset) the spine clusters (∼ 6 to 8 spines) that were stimulated. Scale bars, 5 µm. **(C-E)** The dV/dt of cluster-evoked responses as a function of the response amplitude, distance from the soma, and RMP. Red dots denote Na^+^ spikelet dominated responses. N = 208 clusters from 66 cells in (C) and (D). N=51 clusters from 20 cells in (E). **(F, i)** The standard deviation of uncaging-evoked spike timing (jitter) decreases with strong clusters (Na^+^ spikelet driven plateau potential) in comparison to weak clusters (plateau potential without observable Na^+^ spikelet) (Mann–Whitney–Wilcoxon test, N = 11 strong clusters and 16 weak clusters from 20 cells, p < 0.001). **(F, ii)** Uncaging evoked spike onset delay decreases with strong clusters (Mann–Whitney–Wilcoxon test, N = 14 strong branches and 28 weak branches from 20 cells, p < 0.001). **(G)** Schematic depicting the experimental arrangement in which a set of distributed (across multiple branches) and clustered spines (across the same branch) are holographically stimulated. **(H)** Holographic uncaging was performed across two clusters, one reflecting a Na^+^ spikelet followed by a NMDAR dependent plateau (A, red), one with only a NMDAR plateau (B, green), and a set of distributed synaptic inputs with a somatic depolarization of a few milli-volts (bkg, blue). **(I)** The somatic spiking response (10 repetitive trials each) upon holographic activation of A+bkg and B+bkg. **(J)** Spikelet-mediated temporal precision response under strong and weak cluster activation. **(J, i)** Strong clusters lead to less spike jitter when paired with background synapses (Mann–Whitney–Wilcoxon test, N = 8 strong clusters and 5 weak clusters from 14 cells, p < 0.005). **(J, ii)** Temporal jitter as a function of the Na^+^ spikelet amplitude. **(J, iii)** Two example clusters with large (red) and small (blue) Na^+^ spikelet amplitudes, and the somatic spikes evoked when each cluster is paired with background synaptic inputs. **(K, i)** Schematic of the experimental arrangement showing holographic co-activation of 3 synaptic clusters across multiple different branches. **(K, ii)** An example cell showing the extent of spread of synaptic stimulation (color-coded by z-depth). Scale bar, 20 µm (main) and 2 µm (inset). **(K, iii)** 10 trials under holographic stimulation of up to 3 clusters across 2 different cells showing the AP response when at least one cluster exhibits a Na^+^ spikelet-mediated plateau (left) and when all 3 clusters exhibit a plateau without a Na^+^ spikelet (right). **(L, i)** Schematic depicting the experimental arrangement in which clusters are evoked on different basal dendrites with a distributed background synaptic drive. (right, inset) clusters stimulated. Scale bars, 5 µm. **(L, ii)** Triplets and doublets evoked when clusters paired with background. (**L, iii**) Area under curve of the uncaging waveform (measured at -65mV) for the clusters that generate bursts compared to single APs (Mann–Whitney–Wilcoxon test, N = 15 branches evoking bursts and 41 branches evoking single AP from 20 cells, p < 0.001).

Critically, the change in RMP was enabled by current injection via the patch electrode. We surmised that such a change in RMP could be elicited *in vivo* by relatively slow and weak synaptic background activity. What would be the nature of synaptic patterns needed to mimic sufficient changes in RMP for biasing AP dynamics? Under such conditions, how would multiple clusters interact within and across basal dendrites to cooperatively dictate AP dynamics? To unravel this relationship, we first performed 3D holographic uncaging across two synaptic clusters located along the same basal dendrite while simultaneously stimulating a distributed synaptic input pattern across multiple dendritic arbors (**Figure 2G**). The neuron at rest fluctuated between an RMP of ∼-65mV to ∼ -62mV and no current was injected to maintain this RMP. While one strong cluster (denoted A) showed a distinct Na^+^ spike driven plateau, a second weak cluster revealed a plateau reflective of an NMDAR event but without the Na^+^ component (denoted B) (**Figure 2H**). The distributed input (denoted bkg for background) resulted in a small amplitude (∼ 2 mV to 5 mV) yet slowly varying depolarization which was not sufficient to drive somatic spiking. While cluster A paired with the distributed background resulted in precise AP generation, cluster B paired with the distributed background resulted in a highly jittered AP response (**Figure 2I**), similar to when the RMP was modulated by current injection via the patch pipette. Significantly, and in line with the results from **Figure 2C**, both low and high amplitude Na^+^ driven d-spikes resulted in millisecond AP timing (**Figure 2J (i)**) (Mann–Whitney–Wilcoxon test, N = 8 strong branches and 5 weak branches from 13 cells, p < 0.002) suggesting amplitude of the d-spike (**Figure 2J (ii))** wasn’t a critical determining factor in ensuring precision, but rather the dV/dt (**Figure 2J (iii)**). Importantly, we found that even small dV/dt values (∼2V/s) were sufficient to generate precise spikes, in agreement with earlier reports from CA1 cells (Golding and Spruston, 1998; Losonczy et al., 2008). To evaluate if increasing the number of clusters across dendrites interfered with spike precision generation, we stimulated clusters across multiple basal dendrites separated by at least 20µm (**Figure 2K (I, ii)**). In a set of cells in which at least one cluster reflected a strong Na^+^ component, spike precision was maintained (**Figure 2K (iii), left** jitter = 1.91 +/- 2.28 ms, N = 4 neurons). When none of the clusters reflected a Na^+^ spike but instead reflected large dendritic depolarizations including NMDAR spikes, we observed a jittered response (**Figure 2K (iii), right,** jitter = 7.83 +/- 8.53 ms, N = 2 neurons).

Importantly, when two clusters on separate basal dendritic branches exhibiting Na^+^ spikes were paired with a weak background (**Figure 2L (i)**), the area under the curve (duration of the plateau), correlated positively with the propensity of bursting (**Figure 2L (ii)**). This result showed that dV/dt alone was not critical for bursting but rather the duration of the plateau was a key determining factor (Mann–Whitney–Wilcoxon test, N = 15 d-spike clusters evoking bursts and 41 d-spike clusters evoking single AP from 20 cells, p < 0.001). Such high-frequency bursts were observed to efficiently back-propagate into the basal dendrites compared to a single AP (**Figure S3**). Once again, notably, in the presence of the dendritic Na^+^ spike, the somatic AP output was precise in terms of timing and exhibited no observable jitter. Together, the above results highlight four critical features. First, dendritic Na^+^ spikes potently control somatic spike precision irrespective of background inputs. Second, NMDAR spikes lacking the Na^+^ component but large enough to depolarize the soma above threshold results in a variable spike onset. Third, the duration of the dendritic plateau is critical in determining burst propensity. Finally, in a multibranch framework, a weak synaptic background of just a few milli-volts, not strong enough to evoke an AP in isolation, can still critically bias the axo-somatic region to an optimal condition conducive for precise AP output. This is particularly important as nonlinear voltage-gated conductances in the axo-somatic segment might be regulated by this small yet sufficient background depolarization.

### Persistent axo-somatic Na^+^ current mediates cluster-evoked burst control

We next ascertained if axo-somatic voltage-gated amplification played a role in modulating spike timing and burst generation. It is now well appreciated that neocortical pyramidal neurons exhibit a subthreshold, slowly inactivating steady state inward “persistent” Na^+^ current (*I*_NaP_), which plays a key role in modulating neuronal rhythmicity (Astman et al., 2006). The primary source of *I*_NaP_ is in the proximal axon initial segment (Astman et al., 2006; Bean, 2007) and hence is uniquely poised to impact AP threshold, synaptic integration patterns, and resultant dendritic nonlinearities. *I*_NaP_ is most active between –65 mV and –35 mV, is TTX-sensitive, and exhibits an e-fold change in current for 4–6 mV shifts in membrane potential – a sharp voltage dependence of activation comparable to the fast-inactivating transient sodium current (*I*_NaT_), which is responsible for axo-somatic AP generation. Although the maximum steady-state *I*_NaP_ current (on the order of ∼100 pA) is much smaller than that of *I*_NaT_, the regenerative nature of the current leads to an enhanced excitation at subthreshold voltages (Bean, 2007). To corroborate this subthreshold voltage-dependent amplification, we used theta stimulation to elicit a short burst of unitary EPSP’s (**see Methods for details**), mimicking coincident synaptic inputs. In line with the previous reports (Hsu et al., 2018), we observed a clear subthreshold voltage-dependent amplification at the soma in response to the EPSP burst, which was abolished by TTX but not NMDAR or Ca^2+^ channel blockers (**Figure S4**), suggesting a sodium channel dependence. However, both *I*_NaP_ and *I*_NaT_ are TTX-sensitive and in order to study the effect of *I*_NaP_ on AP dynamics, *I*_NaP_ needs to be selectively blocked without affecting *I*_NaT_. To achieve this, we turned to dynamic clamp (Vervaeke et al., 2006). To isolate *I*_NaP_, a slow voltage ramp was applied which caused *I*_NaT_ to inactivate - leaving *I*_NaP -_the only sodium-sensitive current component (**Figure 3A**). Steady state persistent Na^+^ channel conductance’s were then calculated and fitted with a Boltzmann function, revealing a mid-point and knee-point at -40mV and -60mV respectively (**Figure 3B**), consistent with previous reports (Astman et al., 2006; Bean, 2007). Together, this suggested a region of maximum sensitivity for *I*_NaP_ between - 60mV to -50mV. To evaluate whether *I*_NaP_ regulates passive properties of the membrane, we estimated the steady-state input resistance that is sensitive to TTX near the AP threshold (**Figure 3C**). A significant increase in input resistance mediated by *I*_NaP_ was observed between -70mV to -60mV, suggesting that background subthreshold synaptic inputs, even just a few mV’s in amplitude, would be boosted by axo-somatic *I*_NaP_ through a combination of active regenerative inward currents and an increased input resistance (Ceballos et al., 2017). This effect was further reflected by impedance and power-spectral analysis (**Figure S5 A-F**), which showed that impedance increased as membrane potential got closer to AP threshold. This membrane-voltage-regulated impedance was strongest at low-frequencies, indicating weak background synaptic inputs could be amplified by *I*_NaP_-mediated shifts in input resistance. This was aptly reflected in the amplitude and noise level of subthreshold EPSPs as a function of slow changes in RMP (**Figure S5D**) in line with computational predictions (Jacobson et al., 2005; Manwani and Koch, 1999; Steinmetz et al., 2000). Using a kinetic model of *I*_NaP_ (**Figure S5G**, **see Methods sec. for details**) and our recordings (**Figure. 3B**) we then either removed or added the persistent Na^+^ channel conductance via dynamic clamp (**Figure. 3D, 3E**). We first verified the accuracy of dynamic clamp with a current ramp stimulus (**Figure. 3E**). Upon removal of *I*_NaP_, the slope of the voltage response near action potential threshold decreased. Similarly, introduction of *I*_NaP_ after TTX addition generated a recovery in slope (**Figure. 3E, right**). We then injected an artificial NMDAR current into a L5 pyramidal and dynamically removed *I*_NaP_ (**Figure 3F, (i)**). As expected the AP response at the soma was highly jittered with NMDAR like current inputs. Upon removing *I*_NaP_ we observed a reduction in output gain, i.e. a decrease in both burst propensity **(Figure 3F, (ii))** and excitability (**Figures S5H and S5I**), but the temporal precision surprisingly improved (**Figure 3F (iii)**) (Wilcoxon signed-rank test, N = 11 neurons, p=0.03 for burst probability; p = 0.002 for jitter). This shows that under physiological conditions, while *I*_NaP_ does control the burst probability and overall neuronal excitability, the temporal jitter is infact worsened due to *I*_NaP_-mediated amplification. To investigate the role of *I*_NaP_ in synaptic integration, we performed holographic uncaging while dynamically introducing and removing *I*_NaP_ conductance (**Figure 3 G-I**). Cluster uncaging was performed close to action potential threshold to unravel the maximum impact of *I*_NaP_. We observed that removing persistent Na^+^ channel conductance reduced the propensity for the cell to burst by reducing cluster-evoked doublets to singlets (**Figure 3G (i) and (ii)**), while adding *I*_NaP_ conductance to cells reflecting a single AP upon cluster stimulation resulted in doublets (**Figure 3G (iii)**). Next, we repeated the above experiment but did not bring the cell to threshold via patch current injection. Instead, we stimulated synaptic clusters alongside a natural background of synaptic inputs spread across the arbor, which resulted in a ∼3-4 mV depolarization from rest (∼-65mV to -62mV) (**Figure 3H (i)**). Uncaging-evoked subthreshold responses revealed that removing *I*_NaP_ conductance did not affect the dendritically evoked Na^+^ spikelet but did slightly reduce the width of the plateau potential (**Figure 3H (ii)**). Critically, subtraction of *I*_NaP_ conductance did reduce the cluster-evoked triplet to a doublet (**Figure 3H (iii)**) without significantly altering the onset of APs. Notably, removing *I*_NaP_ across multiple cells showed a clear effect in controlling burst probability (**Figure 3I**) (N=7 clusters from 5 neurons, p= 0.0082, paired-sample students-t-test). These results suggest that *I*_NaP_ controls gain via modulation of the plateau potential width, which in-turn could impact the duration of the spike after-depolarization (ADP). The gain modulation lasts approximately 50ms after the initiation of background synaptic input (**Figures S5J-L**), indicating that the slowly inactivating nature of *I*_NaP_ helps extend the temporal window for integration – a feature that can be heightened under distributed multibranch excitation. While the persistent subthreshold conductance allow for dynamic gain control, it does not seem to underlie precise timing and low AP jitter.

**Figure 3:**
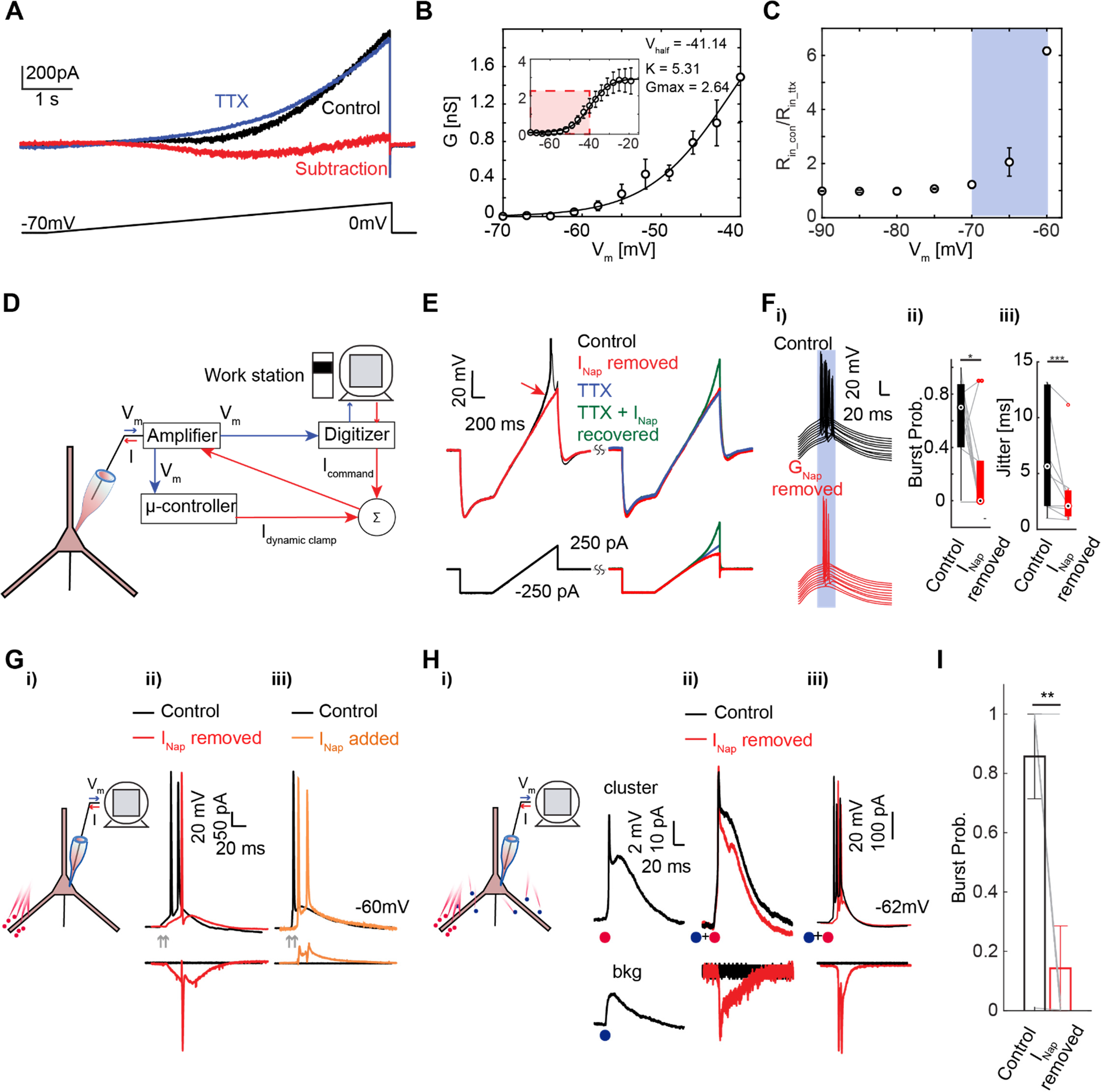
I_NaP_ regulates cluster-evoked high frequency bursting. **(A)** Slow voltage ramp reveals the slow-inactivating (persistent) Na^+^ current. Black: control, blue: 1µM TTX, red: difference between control and TTX conditions. Bottom: voltage ramp at 10mV/s. **(B)** Steady-state conductance estimated based on voltage ramp response from (A) and fitted with Boltzmann sigmoidal function (mean +/- s.e.m, N = 12 neurons). **(C)** Na^+^-current-sensitive rapid increase in input resistance at near-spike-threshold (- 65 to -60mV), estimated with small (5mV) voltage steps (mean +/- s.e.m, N = 8 neurons). **(D)** Schematic depicting the layout of the dynamic clamp setup. **(E)** Representative response of a current ramp under control (black), subtraction of I_NaP_ (red, G_NaP_ = -4 nS), 1µM TTX added to the bath (blue), and recovery of I_NaP_ via dynamic clamp (green, G_NaP_ = 4nS). Current injected to the cell shown below. **(F, i)** Somatic spike response to dynamic clamp injection of a NMDAR-plateau potential response under control (black) and with I_NaP_ removed (red, G_NaP_ = -3nS). **(F, ii and iii)** Spike burst probability and timing jitter under control and conditions in which I_NaP_ is removed (Wilcoxon signed-rank test, N = 11 neurons, p=0.03 for burst probability, p = 0.002 for Jitter). Note the increase in somatic gain but decrease in AP temporal precision with I_NaP_ present. **(G, i)** Schematic of the experimental setup of the dynamic clamp alongside holographic uncaging. **(G, ii and iii)** Dynamic removal of I_NaP_ (red, G_NaP_ ∼ -4nS) reduces a burst to a single spike (shown in red) while dynamic addition of I_NaP_ induces bursts (orange, G_NaP_ = 1.5nS). Note the dramatic reduction in the spike after-depolarization which reduces but does not completely cease. **H-I)** I_NaP_ mediates AP bursts induced by cooperation of synaptic inputs. **(H, i)** Somatic response evoked by uncaging synaptic clusters (red dot) and a background of distributed inputs (blue dot). **(H, ii and iii)** Coactivation of the cluster and distributed background under control (black) and with I_NaP_ removed (red, G_NaP_ ∼ -3nS). Corresponding current command is shown below. Note that removal of the persistent Na^+^ current did not change the dendritic Na^+^ spikelet evoked plateau potential waveform. **(I)** Somatic burst probability under control (black) and with I_NaP_ removed (red) (N = 7 clusters from 5 neurons, p = 0.0082, paired-sample student-t-test).

### Dendritic Na^+^ spike driven stochastic channel gating explains somatic AP precision

Persistent Na^+^ currents modulate somatic AP gain and suppressing them lead to increased jitter. A different mechanism must therefore exist to ensure precise and jitter free spike initiation, especially when dendritic Na^+^ spike-driven nonlinearities drive somatic output (Ariav et al., 2003). Previous experimental reports have suggested that precise APs occur when the stimulus is dynamic, time-varying, and noisy instead of quasi-static (Mainen and Sejnowski, 1995). However, models of spike initiation and propagation often fail to capture the dynamic nature of dendritic voltage-gated Na^+^ channels and their role in initiating or augmenting a somatic AP. There is, however, growing evidence that the probabilistic modulation of voltage-gated channels infuses noise into the trans-membrane current and is a key determining factor in spike reliability. In particular, the frequency content of the noisy stimulus impacts spike precision (White et al., 2000). Since dendritic Na^+^ spikes reflect a fast fluctuation with high-frequency content, they could serve as an ideal substrate to gate axo-somatic channels in a stochastic manner, enabling efficient AP generation. To reveal the role of stochastic gating by dendritic Na^+^ spikes, we resorted to a combined experimental-computational framework. First, just as in the seminal study by Mainen and Sejnowski (Mainen and Sejnowski, 1995), we stimulated the cell with pseudorandom noise superimposed on a DC current background and found that spike onset jitter reduced with decreasing time constants (τ) of injected current transients (**Figure 4A (i, ii)** Wilcoxon signed-rank test, N = 13 neurons, one asterisk: p < 0.05, two: p < 0.01, three: p < 0.005). The dendritic Na^+^ spike, given the fast kinetics of Na^+^ channels (Ariav et al., 2003), has a tau on the order of a millisecond (**shaded window in Figure 4A (ii)**), which may explain the corresponding high temporal precision of APs observed. This would suggest that instead of average input amplitudes, the fast fluctuation of the dendritic spike might be responsible for optimal channel gating. To investigate the possible mechanisms that enable fast fluctuations (steep current slopes) to enhance temporal precision, we numerically simulated the probabilistic description of Hodgkin-Huxley style Na^+^ and K^+^ channel activities. Modeling gating action of voltage-dependent ion channels using stochastic state transitions (Chow and White, 1996; Schneidman et al., 1998) has been proposed to explain spike timing reliability and precision, as a function of fluctuating pseudorandom current inputs. Following these studies, we used a 13-state Markov Chain description of Hodgkin-Huxley-style ion channels (voltage-gated Na^+^, delayed rectifying K^+^, and leak) incorporated with the stochastic theory (**see method section, Figure S6A, S6B**) and reproduced the experimental results of **Figure 4A**, in which fast dynamic currents evoked APs with high precision (**Figure S6 C and D**). To mimic the d-spike-induced AP output with this model, we injected our experimentally measured cluster-evoked dendritic voltage transients, with and without dendritic Na^+^ spikes, as currents to the 13-state stochastic neuron model. Na^+^ spikelet dominated dendritic nonlinearities evoked high precision somatic APs (**Figure 4B (i), top**) while NMDAR events alone resulted in an appreciable jitter (**Figure 4B (i), bottom**). These results verify that the time course of dendritic Na^+^ spike is sufficient for enhancing spike precision via stochastic gating.

**Figure 4:**
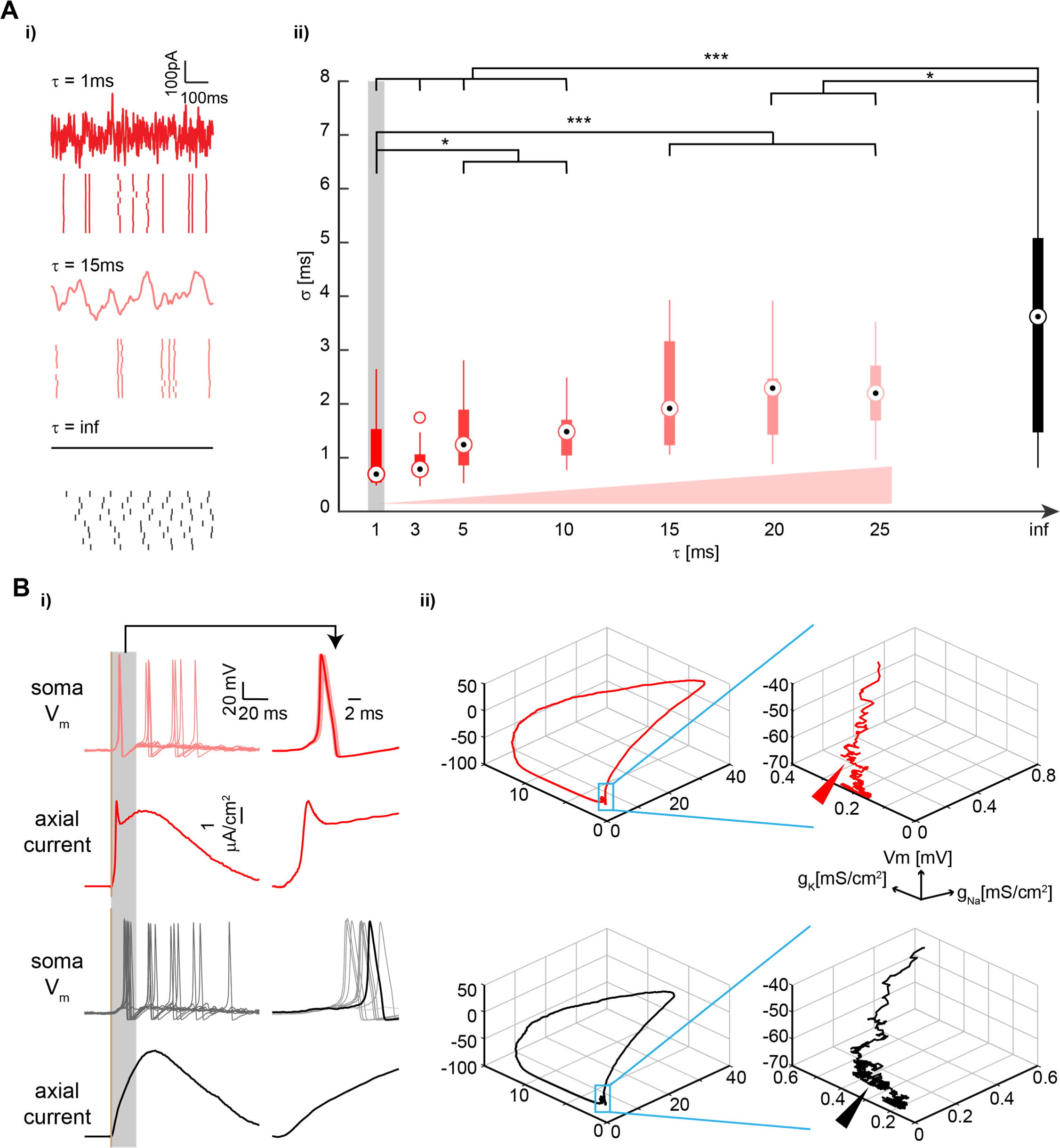
Stochastic gating regulates the spike temporal precision. **(A, i)** Representative pseudo-random noisy current injections with various time constants τ (top), and corresponding spiking raster of 10 repetitive trials (bottom). **(A, ii)** Spike jitter decreases with fast fluctuations of input stimulus. Shaded window represents likely time constant of Na^+^ spikelet-driven responses. (Wilcoxon signed-rank test, N = 13 neurons, one asterisk: p < 0.05, two: p < 0.01, three: p < 0.005.) **(B, i)** (Top) Membrane potential response of the stochastic HH model (10 repetitive trials) under current injection with a Na^+^ spikelet driven d-spike(red) or NMDAR d-spike (black) waveform. (Bottom) injected current. The current waveform was experimentally extracted from uncaging experiments, smoothed and the amplitude adjusted to 6 µA/cm^2^ to match with the excitability of HH model. **(B, ii)** Phase plane trajectory of Na^+^ and K^+^ channel dynamics under the conditions in (B, i). (Inset) Spike initiation period. Arrows: Spontaneous ion channels gating during the spike initiation. Note the larger variability in the transition dynamics of ion channel gating towards spike initiation.

Mechanistically, the reason for why rapidly varying input currents increase spike precision may be the combination of the highly bi-stable nature of action potential generation and rapid activation and inactivation of a small number of transient Na^+^ channels (**Figure 4B (ii)**) (Schneidman et al., 1998). Specifically, the interaction of ion channels can be approximated to a dynamical system with resting (i.e. a stable basin of attraction) and firing states. Action potential generation near threshold is highly non-linear, wherein a relatively small perturbation would push ion channels into a “feed-forward” phase, transitioning from the resting to firing state. In other words, a small number of Na^+^ channel openings activated simultaneously would be sufficient to initiate an action potential. Since a majority of axo-somatic Na^+^ channels are in a resting state below threshold, a sudden rapidly varying input driven by Na^+^ dendritic spikes would “push” a sufficient number of axo-somatic Na^+^ channels to activate in a short time window with a straight-forward trajectory in phase space (**Figure 4B (ii), red trace, inset with arrow**). Since these channels have very little time to inactivate, the transition to a firing state is optimal and precise. On the contrary, under a slowly varying input driven by NMDAR spikes the Na^+^ channels would activate but also inactivate (**Figure 4B (ii), black trace, inset with arrow**). This would ensure that across repetitive identical inputs, the exact time at which a sufficient number of Na^+^ channels will open becomes variable. Significantly, spike precision isn’t merely the result of current injection above threshold, but a consequence of the interaction between ion channels and input current at/near the bi-stable point. This would explain why suprathreshold synaptic input won’t lead to optimal AP precision (**Figure 4A**). This result also highlights why dendritic Na^+^ spikes which exhibit a large dV/dt but not necessarily large amplitudes are sufficient to drive an optimal number of axo-somatic Na^+^ channels into the firing state.

### Clustered synaptic stimulation evoked precision and gain control under noisy *in-vivo* like states

The results presented in **Figure 2** and the associated mechanisms described in **Figures 3 and 4** were all performed under *in vitro* quiescent conditions. We set out to confirm if AP precision and gain modulation as a function of clustered synaptic stimulation and multibranch integration held under *in vivo*-like states with a noisy fluctuating background and high-conductance. First, we corroborated spike timing statistics and background firing rates under feedforward sensory sweeps in awake head-fixed mice. This was needed to recapitulate the background firing rate range *in vitro*. Mice were habituated to a wheel and high-density silicon probes were inserted into the C1 column of the barrel cortex (localized via intrinsic imaging). Upon contralateral somatotopically aligned whisker stimulation (**Figure 5A**) we verified the cortical layers probed via depth of probe insertion tracks (**Figure S7A**) and current source density analysis (**Figure 5B**). Single unit responses time-locked to whisker stimulation were observed in both L2/3 and L5 in the form of both isolated units and high-frequency bursts (**Figure 5C**). The peri-stimulus time histogram (PSTH) of representative single units showed a highly time-locked peak within 20ms of whisker deflection (**Figures 5C and 5D (i)**). A majority of L2/3 and L5 single units showed a time-locked peak in PSTH (**Figure S7B,** n = 32 L2/3 single units and 137 L5 single units from 6 mice). The average firing rate of L5 stimulus-evoked units in the barrel cortex of awake mice was below 10Hz, with a majority displaying a firing rate between 2 to 6 Hz (**Figure 5D (ii)**). Between 20% to 40% of the measured responses, however, comprised of bursts with frequencies of 50Hz and higher (**Figure S7C,** n = 137 L5 single units from 6 mice). Under such conditions of background firing and high-conductance conditions *in vivo* (Destexhe *et al*., 2001), precisely how coincident inputs in a multibranch framework integrate to control AP precision and gain is relatively poorly mapped.

**Figure 5:**
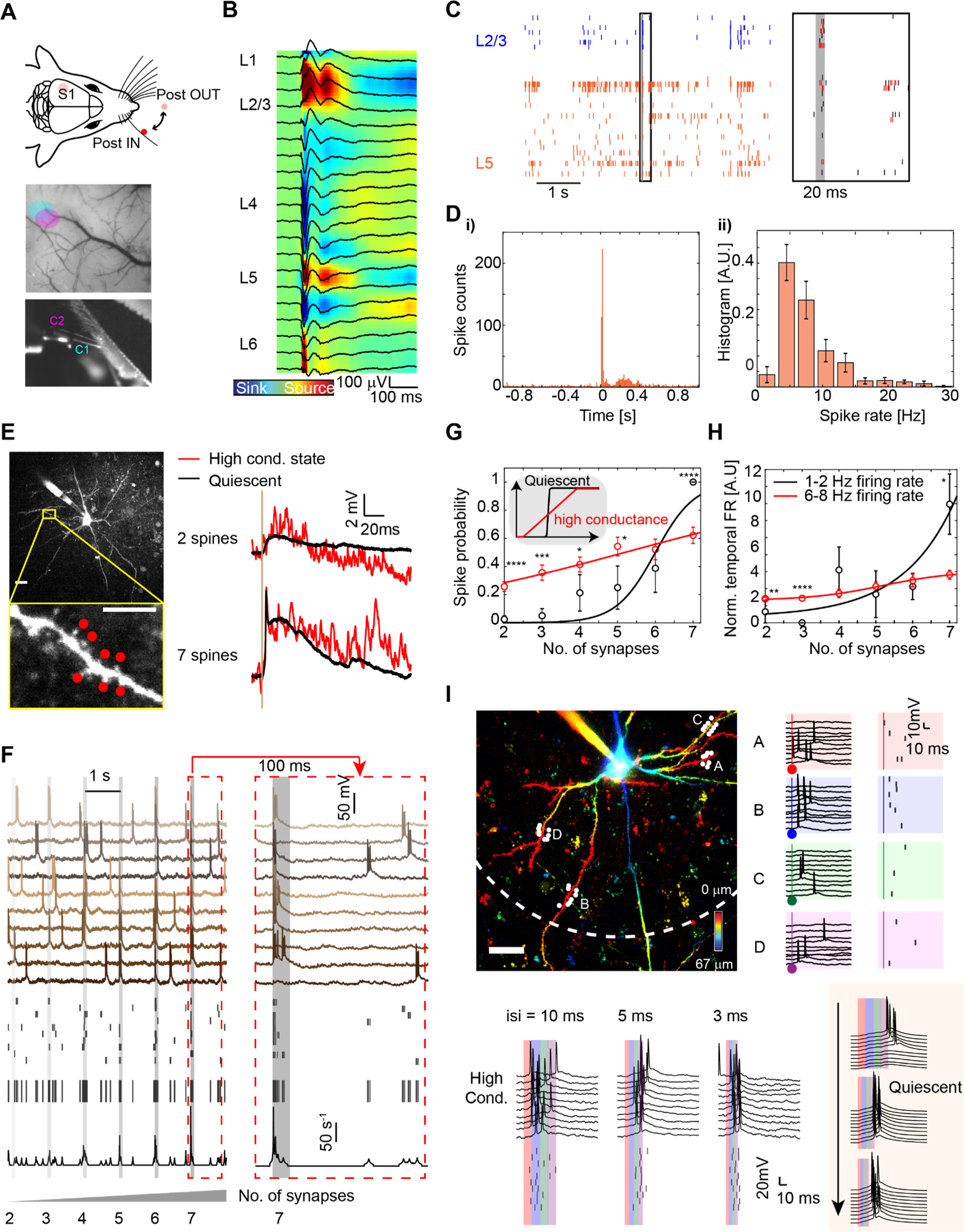
Uncaging-evoked spike timing and gain modulation under conditions of high background noise and conductance. **(A)** (Top) experimental setup to map sensory evoked responses in the awake barrel cortex. (Bottom) intrinsic imaging response to whisker stimulation for silicon probe insertion. **(B)** Average Local field potential (LFP) and current source density (CSD) upon whisker deflection (interelectrode spacing, 50 µm). **(C)** Whisker touch-evoked single-unit raster’s in the C1 barrel cortex of awake mice. (see text and methods for details). Both singlets (black) and bursts (red) are observed. **(D, i)** Peri-stimulus time histogram (PSTH) of a representative single unit in L5 reveals precise spike onset upon whisker stimulation. **(D, ii)** Histogram of average firing rate in L5 (mean +/- s.e.m, N = 6 animals). **(E)** Examples of cluster-evoked uncaging response under dynamic conductance injection. Scale bars, 20µm, 5µm (inset). Note that even under high-excitatory drive and noisy backgrounds the uncaging response is clearly observed. **(F)** Na^+^ d-spike mediated spike precision is maintained under high-background noise and conductance conditions by clustered synaptic activation. Holographic excitation of spines ranging from 2 to 7 in number across 10 trials clearly showed that temporal firing rate (bottom, black) was augmented by clustered inputs. **(G)** Background noise and increased conductance improves probabilistic computation. (Mann–Whitney–Wilcoxon test, N = 14 clusters from 8 neurons, one asterisk: p < 0.05, two: p < 0.01, three: p < 0.005, four: p < 0.001) (Inset, top) theoretically expected trend in spike probability as a function of synaptic drive under low and high conductance state. (**H)** Clustered synaptic inputs increase relative temporal firing rates with ongoing background firing. (Mann–Whitney–Wilcoxon test, N = 14 clusters from 8 neurons, one asterisk: p < 0.05, two: p < 0.01, three: p < 0.005, four: p < 0.001). (**I)** High conductance-like-states enhance temporal resolution during multibranch integration. (Top, left) two-photon depth encoded image showing an example of 4 clusters (7 to 8 spines each cluster) stimulated synchronously across basal dendritic arbors. Dashed arc marks a 100µm radius from the soma (Top, right) uncaging evoked response across multiple trials for each cluster. (Bottom, left) Holographic stimulation of all 4 clusters under the high-conductance-like state showing increased spike generation probability, low onset delay, and fast responsivity as a function of inter-cluster delay. (Bottom, right) Same response under quiescent conditions indicates spike generation only after all four clusters have integrated. (N=60 trials from 2 cells, p < .0001, Mann–Whitney–Wilcoxon test; (not shown)) (Scale bars, 20 µm)

The *in vivo* like high conductance state (Destexhe et al., 2003) is characterized by low input resistance, noisy membrane fluctuations, a highly depolarized RMP, and a significant amount of background AP firing. Traditionally, high-conductance states are modeled using a near-equal inhibitory and excitatory conductance (Fernandez et al., 2011). We however, maintained a static high-excitatory drive (2nS to 4nS) with a moderate inhibitory drive (1 nS to 2 nS) to avoid extreme shunting while still being able to elicit a response from 6-10 spines per cluster. Under such conditions, noisy membrane fluctuations resembling *in vivo* recordings (Jayant et al., 2019) (2 to 4 mV rms), a decrease in input resistance (mean R_in_ = 59 ± 14 MΩ) consistent with the range obtained from whole-cell recordings from L5 neurons in the awake S1 (Zhao et al., 2016), and increased background firing rates, matching our *in vivo* recordings (**Figure 5D (ii)**) and in line with recent reports (de Kock et al., 2021), were observed. We created a high conductance dynamic input based on these parameters (**see Methods for details**), while holographic uncaging was performed across clustered synapses spanning a single dendrite (**Figure 5E and 5F**). We found that under such conditions the characteristic uncaging-evoked dendritic Na^+^ spike-driven plateau was preserved (**Figure 5E**). Importantly, stimulation of just a few spines resulted in a noisy yet clearly discernible EPSP at the soma.

One could question the validity of performing a dynamic clamp at a somatic point source for influencing uncaging-evoked input-output transformations. To justify our approach, the conductance visibility observed from the soma for a passive cable is well approximated by λ/2 (Koch et al., 1990; Williams, 2004), where λ is the steady-state space constant of basal dendrites (∼250µm) (Nevian et al., 2007), and this length approximates to ∼120 µm from the soma for L5 pyramidal neurons (**Figure S8, and Supplemental Note 1**). Moreover, under cable like properties and given the mismatch in input resistances between the soma and basal dendrites, synaptic conductance generated across basal arbors is also expected to be visible at the soma. Therefore, a dynamic conductance injected at the soma robustly reflects widespread basal dendritic drive (Koch et al., 1990; Williams, 2004). Given that dendritic conductance visibility spans ∼100um, it serves as a reasonable approximation for elucidating the effect of high conductance on synaptic integration. We stimulated individual spines and clusters across a single dendrite in sequence and observed that despite the background noise and ongoing voltage-resets that might be caused by back-propagating APs, uncaging-evoked Na^+^ d-spike-related temporal precision was maintained (**Figure 5F**; jitter = 3.16 +/- 4.11 ms, N = 5 clusters from 4 neurons). Varying the number of spines within the cluster showed that clustered stimulation of ∼6 to 7 spines clearly resulted in a precise AP output. Notably however, in some trials, co-activation of only 2 or 3 synapses within the cluster would result in a somatic spike, unmasking an effect wherein a weak synaptic drive that was usually insufficient to modulate the somatic output under quiescent conditions now had a finite probability to generate a somatic AP under the high conductance state (**Figure 5G**) (Mann–Whitney–Wilcoxon test, N = 14 clusters from 8 neurons, one asterisk: p < 0.05, two: p < 0.01, three: p < 0.005, four: p < 0.001). Significantly, this result shows an improved dynamic range at the level of individual synapses enabling probabilistic gating of AP output under a high conductance state. This result validates computational predictions put forth by Destexhe and colleagues (Destexhe et al., 2003; Hô and Destexhe, 2000) providing the first experimental demonstration of synaptic resolution efficacy on spike generation under noisy backgrounds. Strikingly, increasing the background firing rates up to 10Hz did not saturate the effect of uncaging-evoked responses indicating that voltage-resets due to back-propagating APs is not detrimental to coincident input efficacy (**Figure 5H**) (Mann–Whitney– Wilcoxon test, N = 14 clusters from 8 neurons, one asterisk: p < 0.05, two: p < 0.01, three: p < 0.005, four: p < 0.001). Together, the results shown in **Figures 5G and 5H** suggest 1) background noise enhances the ability of weak inputs to initiate an AP; 2) clustered synaptic stimulation ensures somatic spike precision even in the presence of noisy fluctuations and ongoing background activity 3) back propagating action potentials caused minimal somato-dendritic cross talk to disrupt clustered synaptic integration.

Dendritic cross-talk is most severe under quiescent conditions (Behabadi and Mel, 2014) as the voltage emanating at the soma would be felt across a vast section of the basal dendritic arbors. This is intensified by a steady state length constants of ∼450 µm (Nevian et al., 2007) in L5 pyramidal neurons at rest. This effect would be different under high conductance conditions as the somatic depolarization for unitary inputs a) would be reduced due to lower input resistance; b) would decay faster in the basal dendrites due to the reduced time constant. To examine the nature of cooperativity and the efficacy of clusters in modulating somatic spike timing and gain under the high-conductance-like states we stimulated four synaptic clusters across multiple basal dendrites (∼ 50 µm to 80 µm from the soma) from near simultaneous to sequential with a delay set by shifts in phase mask of the SLM. Under dynamic clamp the increased conductance at the soma should filter back into the basal dendrites over ∼120µm. Under these conditions, we observed that irrespective of the delay in between cluster stimulation, somatic AP responsivity and gain was enhanced by synchronous stimulation across branches (**Figure 5I**). Notably, no single cluster saturated or disrupted the spike generating ability of another cluster. Infact, when multiple clusters were stimulated near synchronously, the gain, and responsivity were enhanced reflecting a more efficient integration and temporal firing rate enhancement, including across multiple trials (N = 60 trials from 2 cells, spike onset delay = 20.2 +/- 14.3 ms under the high-conductance-like state, 32 +/- 12.1ms under the quiescent conditions, p < 0.0001, Mann– Whitney–Wilcoxon test). On the contrary, under quiescent conditions, all four clusters had to be stimulated to generate a reliable spike, indicating a much slower integration and lower responsivity – an effect possibly exacerbated by increased membrane resistance and tau. Such efficient integration under increased-conductance drive and noisy fluctuations could be a critical way for the neuron to add spike modules to ongoing activity *in vivo*.

## DISCUSSION

In the present study, we customized, characterized, and performed 3D two-photon SLM-based holographic transmitter uncaging across multiple basal dendritic arbors of L5 pyramidal neurons under quiescent and *in vivo* like conditions. We combined holographic uncaging with whole-cell electrophysiology, dynamic clamp, and biophysical modeling and examined how co-active dendritic nonlinearities across multiple branches multiplex at the soma to control somatic spike timing and gain. Specifically, we investigated the intrinsic mechanisms underlying precise AP timing and burst control under synchronous synaptic streams (clustered and distributed). We delved into the role of stochastic channel gating in modulating AP timing and axo-somatic persistent sodium currents in modulating AP gain. Finally, we explored the role of synaptic clusters in enabling temporal precision and gain under *in vivo* like conditions, and queried the extent to which just a few synapses and clusters could impact somatic spiking under noisy background conditions.

Basal dendrites are known to exhibit a rich repertoire of regenerative dendritic spikes mediated via Na^+^ and NMDAR channels. These regenerative events are a strong function of synchronous synaptic inputs and there is mounting evidence that such synapses are clustered across a 5 µm to 10 µm length scale (Gökçe et al., 2016; Scholl et al., 2021; Scholl et al., 2017). In the cortical slice preparation, a vast majority of L5 basal dendrites radiate outwards spanning several tens of microns. Hence, probing multisite multibranch dendritic integration using traditional galvanometric uncaging is not feasible. Our results clearly show that 3D two-photon holographic uncaging enables clustered synaptic stimulation with single-spine resolution spread across multiple basal arbors spanning a depth of ∼40 µm – 70 µm. Using this innovative methodology, we demonstrate multi-cluster stimulation in sequence across multiple arbors with no observable photodamage or excitotoxicity. Almost all of our current understanding of clustered synaptic stimulation has come from uncaging evoked responses across short dendritic segments using rapidly switching galvanometric mirrors. The ability to stimulate clusters of synapses at will across the entire length of the dendrite, as well as across multiple dendrites, opens up the possibility of querying synaptic and dendritic integration, cooperativity, and spike-generation mechanisms in much greater detail.

Clustered synapses have been envisaged to underlie feature binding, memory storage, and pattern recognition. This would imply that clustered synapses should be able to modulate the firing properties of the neuron in a meaningful way, for example, via dynamic changes in threshold (Azouz and Gray, 2000). Signaling of such synchronized inputs in the presence of voltage-gated channels, ongoing background activity, and noise – both in the form of stochastic membrane potential fluctuations as well as a background rate code, is critical for efficient neural computation (Silver, 2010). We investigated the mechanisms through which such clustered activation across single as well as multiple dendritic segments impact action potential generation. Our findings (**Figure 2**) demonstrate that non-linear Na^+^- mediated dendritic spikes drive action potential activity with high-precision, while NMDAR plateau potentials mediate gain control. Importantly, these nonlinearities modulate voltage-gated axo-somatic persistent Na^+^ channels to control output gain (**Figure 3**). Notably, just a few mV’s of depolarization from rest invoked an inward current of ∼200pA (e-fold change in current) which resulted in a nonlinear increase in input resistance between -70mV to -60mV, which, being close to threshold could play a key role in amplifying weak inputs (**Figure 3C**). Moreover, we found that this persistent Na^+^ channel-dependent gain control was robust over an appreciable time window of ∼50ms indicating a large window of integration that can be opened by background synaptic activity – an effect that could be critical for spike-timing-dependent plasticity (Duguid and Sjöström, 2006; Legenstein and Maass, 2011; Tazerart et al., 2020). Nonlinear integration across the basal dendritic tree in pyramidal neurons follows the well-corroborated augmented 2-layer model of computation (Jadi et al., 2014) wherein a sequence of 2D sigmoidal input-output transformations across the dendritic tree accurately reflects spatial integration effects. This 2D sigmoidal property provides pyramidal neurons with an extended range of nonlinear computational capabilities. Nonlinear modulation by persistent Na^+^ currents at the axo-somatic junction by dendritic spikes could further improve this 2D augmented model of integration. Somatic spike precision improved upon removing persistent Na^+^ channels (via dynamic clamp) and was not dependent on the amplitude of plateau potentials, but instead dependent on the sharp fluctuations in voltage elicited by dendritic Na^+^ spikes. A model based on stochastic gating of channel activation near threshold captured this phenomenon (**Figure 4**), suggesting that sharp voltage fluctuations from dendritic nonlinearities could play a critical role in creating an energetically favorable mode of spike generation. – i.e. the activation of just a few channels near threshold can push the neuron from state of rest into a state of firing. The probability of a few channels transitioning at the same time is critically dependent on the variance in the signal, which is appreciable in the case of dendritic Na^+^ spikes. Since dendritic Na^+^ spikes have been observed to occur at high rates *in vivo* (Destexhe and Mehta, 2022), it could imply that stochastic gating is critical for energy efficient computations (Schneidman et al., 1998; White et al., 2000).

Neurons *in vivo* are however characterized by noisy membrane fluctuations, low input resistance, and depolarized RMPs. We examined how clustered synaptic inputs, which under quiescent conditions results in dendritic nonlinearities, impacts AP generation under *in vivo*-like conditions. We found that precision and rate control are maintained under conditions of low input resistance and high background noise. Computational efforts examining integration amidst such conditions have revealed two key principles (Destexhe et al., 2003), namely, gain modulation via conductance noise, and increased temporal resolution. Our results (**Figure 5**) show that the somatic AP response is enhanced under noisy backgrounds. Specifically, under conditions of increased conductance and noisy membrane-voltage fluctuations, the responsiveness of pyramidal neurons to synaptic stimulation improved with just a few spines (as few as 2 spines) able to elicit a somatic spiking response (**Figure 5F, 5G**), a facet contrary to previous reports (Polsky et al., 2009). Such sparse inputs however, did not elicit somatic spiking under quiescent conditions. This result suggests that sparse inputs and their interaction with a noisy background might be capable of probabilistically modulating somatic spiking, especially *in vivo*, which is in line with previous theoretical predictions (Destexhe et al., 2003; Goetz et al., 2021). Clustered synaptic inputs, in contrast, had a profound impact on AP generation ensuring precision, gain, and increased temporal resolution consistently across multiple dendrites (**Figure 5H, 5I**). This increased temporal responsivity of AP generation agrees well with the lower overall time constant of the neuron under high-conductance conditions. While the lower voltage drive and reduced input resistance under high-conductance conditions would lead to a reduced amplitude of dendritic nonlinearities and in turn a lower somatic gain, clustered synaptic inputs could help maintain efficient spiking output amidst ongoing noise (Scholl et al., 2021). This view complements a recent study in which it was shown that even a few synapses can recruit sufficient NMDAR nonlinearities in basal dendrites due to the low AMPA:NMDAR ratio (Lafourcade et al., 2022). This would suggest that NMDAR currents along with persistent Na^+^ currents, which would already be active due to the highly depolarized RMP, would open a large window of integration to enable AP gain control.

We also observed that such increased AP responsiveness at the soma was not affected by backpropagating action potentials. This is crucial because during backpropagating APs, a vast stretch of basal arborization (∼200 µm) experiences a synchronous voltage reset due to invading transients (Antic, 2003) and this action could impair synaptic and dendritic integration. Computational studies examining the underlying mechanisms of dendritic autonomy amidst such a background barrage allude to the time-course of APs in comparison to NMDAR-mediated nonlinearities (Behabadi and Mel, 2014), which would under conditions of high-conductance equilibrate rapidly in the dendrite and cause minimal voltage-resets. The ability to generate precise APs and burst patterns amidst noise and ongoing backpropagating voltage-resets reasserts that stochastic gating, gain modulation via dendritic nonlinearities, and subthreshold current amplification play a critical role in seamlessly interweaving precision and rate coding, thereby enabling neurons to use both coding strategies in parallel (Destexhe et al., 2003; Tiesinga et al., 2008).

Taking the results in **Figures 3, 4, and 5** we suggest a model for multibranch dendritic integration and AP generation. We propose that widespread isolated background synaptic inputs modulate and tune the gain of the neuron via activation of slowly inactivating sub-threshold voltage-gated channels, while maintaining precision and burst control via dendritic Na^+^ and NMDAR spikes respectively. Importantly, the degree of subthreshold channel activation and strength of dendritic depolarizations results in singlets, doublets and triplets. Overall, this mechanism leads to a scenario wherein feedforward sweeps to the basal arbors can be encoded at the soma via frequency-division-multiplexing. The impact of such a pattern on apical dendritic processing can be profound, especially in reducing the threshold needed for invoking calcium bursts (Larkum et al., 1999) (unpublished observations). Importantly, the proposed mechanisms could help explain how neurons in the awake brain perform feature binding (Costa and Sjöström, 2011; Legenstein and Maass, 2011) despite increases in ongoing firing rates.

## SUPPLEMENTAL INFORMATION

Supplemental information accompanies this article.

## ACKNOWLEDGMENTS

We thank all members of the NanoNeurotechnology lab for critical feedback. We thank Daniel Gonzales for help with intrinsic imaging. We thank Tamara Kinzer-Ursem for sharing mice with us during the early days of the project. This project was supported in part by the R21 EB029740 (NIH Trailblazer); HFSP RGY0069; Ralph E. Powe Junior Faculty Enhancement Award to K.J; and the Purdue Institute for Integrative Neuroscience.

## AUTHOR CONTRIBUTIONS

S.X., S.Y., and K.J. designed experiments. S.X. performed *in vitro* and *in vivo* experiments, characterized the SLM microscope, analyzed the data, wrote code for modeling, and made figures. S.Y. optimized and characterized the SLM microscope with table top compression optics, performed detailed diagnostics, assisted with critical uncaging experiments and figures, and analyzed the measures. S.X., S.Y., wrote the original draft. K.J. conceived and supervised the overall study, and wrote the final manuscript.

## RESOURCE AVAILABILITY

### Lead contact

Correspondence and requests for materials should be addressed to Dr. Krishna Jayant

### Materials availability

This study did not generate any new materials or unique reagents

### Data and code availability

The datasets generated during the current study are available from the corresponding author upon request. All code used in the biophysical models will be made available upon request. Any additional information required to reanalyze the data reported in this paper is available from the lead contact upon request.

## METHOD DETAILS

### Acute slice preparation

All experimental procedures were conducted in accordance with the guidelines set forth by the NIH and Purdue Institutional Animal Care and Use Committee (IACUC). All physiological solutions were fully oxygenated (95% O_2_ and 5% CO_2_), pH adjusted to 7.3 to 7.4, and osmolarity maintained between 300 to 310 mOsm unless stated. Adult C57BL/6 (Jackson Laboratory) mice (both male and female, 6 to 24 weeks of age) were deeply anesthetized with 3 to 4% isoflurane followed by trans-cardiac perfusion with ice-cold NMDG cutting solution (Ting et al., 2014; Ting et al., 2018) consisting of (in mM): 92 NMDG, 30 NaHCO_3_, 1.2 NaH_2_PO_4_, 20 HEPES, 2.5 KCl, 25 glucose, 5 sodium ascorbate, 3 sodium pyruvate, 2 thiourea, 0.5 CaCl_2_, 10 MgCl_2_, 5 N-acetyl-L-cysteine, before decapitation . Coronal slices (300 µm to 350 µm thickness) were prepared using a vibratome (Leica VT1200S) in 0° to 4° Celsius NMDG cutting solution. Brain slices were then allowed to recover in 34°C NMDG cutting solution with gradual spike-in of 0.5 to 1 ml Na^+^ rich NMDG solution (2M NaCl in NMDG cutting solution) over 5 to 10 minutes (time dependent on mouse age) and transferred to room temperature HEPES artificial cerebrospinal fluid (ACSF) holding solution consisting of (in mM): 92 NaCl, 2.5 KCl, 1.25 NaH_2_PO_4_, 30 NaHCO_3_, 20 HEPES, 25 glucose, 5 sodium ascorbate, 3 sodium pyruvate, 2 thiourea, 2 CaCl_2_, 2 MgCl_2_, 5 N-acetyl-L-cysteine for at least 1h prior to recording.

### Electrophysiological recording

Slices were transferred to a chamber, continuously super fused with oxygenated ACSF, and visualized with an upright two-photon microscope (Bruker Nano, Madison, WI) comprising of an Olympus BX51WI body (Olympus, Tokyo, Japan) fitted with infra-red (IR) Dodt-gradient-contrast (DGC) optics, an IR sensitive camera (IR-2000, Dage-MTI, Michigan City, IN), and a 40x water immersion objective (0.8 NA, Nikon USA). Recordings were performed in 34°C recording ACSF consisting of (in mM): 125 NaCl, 3 KCl, 25 NaHCO_3_, 1.25 NaH_2_PO_4_, 25 glucose, 3 sodium pyruvate, 1 sodium ascorbate, 1.3 CaCl_2_, 1 MgCl_2_. 4 to 6 MΩ borosilicate patch pipette (Sutter Instruments, CA, USA) were pulled using a P1000 pipette puller (Sutter Instruments, Novato, CA), filled with internal solution containing (in mM): 130 potassium gluconate, 7 KCl, 10 HEPES, 5 NaCl, 35 sucrose, 2 MgSO_4_, 2 sodium pyruvate, 4 Mg-ATP, 0.4 Tris GTP, 7 phosphocreatine disodium (pH 7.3, osmolarity 290mOsm). 25 µM Alexa 594 or 100 µM Alexa 488 was used for two-photon structural imaging, 200 µM fluo-4 was used for two-photon Ca^2+^ imaging. L5 neurons were patch-clamped and had resting membrane potentials between -60 mV and -70 mV at rest without any current injection. Currents of ∼-50pA to -100 pA was injected in cases where the RMP was more negative (-75mV) or needed to be maintained. Whole-cell recordings were made using a Multiclamp 700B, (Molecular devices, San Jose, CA), Bessel-filtered at 4kHz, and digitized at 4 to 20 kHz using a Digidata 1550B interface (Molecular devices, San Jose, CA) and winwcp software.

### Dynamic clamp

A microcontroller-based circuit board (Desai et al., 2017) was used for performing dynamic clamp experiments. The module was controlled with custom-written codes (Arduino, Processing). First, the linear input-output relationship of the amplification circuit was measured with a DC voltage source (E3631A Agilent Technologies, Santa Clara, CA) and oscilloscope (Keysight, Santa Rosa, CA), and then verified with a model cell (Molecular devices, San Jose, CA). The working principle is as follows: the microcontroller reads the output of the amplifier and then A/D converts the amplified membrane voltage, computes the current based on differential equation models, and D/A converts the current, which is then summed with the current command from the Digidata 1550B interface (Molecular devices, San Jose, CA) (**Figure. 4D**). The ODEs are solved using the forward Euler method, and the time interval (*dt*) for solving the differential equations depended on the processing time of the microcontroller to solve one time point, which ranged from 10 µs to 100 µs, sufficient for neuron electrophysiology with the milli-second computational precision.

A Hodgkin-Huxley style persistent Na channel model (**Eq. 1-3**) was used for dynamically adding or subtracting *I*_NaP_ (*τ* = 1ms, V_half_ = -45.02mV, V_slope_ = 5.01 mV, fitted from experimental results in **Figure. 4B**). m denotes to the activation gating variable of I_Nap_ channels. Given the slow-inactivation timescale of *I*_NaP_ and to avoid a large *dt* due to the processing time of the microcontroller, the inactivation was neglected in this model:

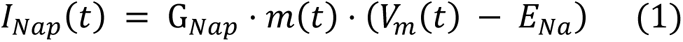

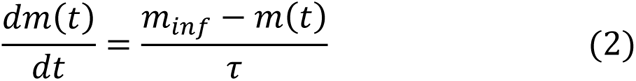

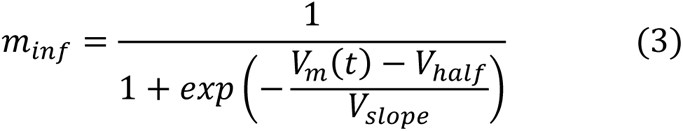

The Ornstein-Uhlenbeck process-based point conductance model (Destexhe et al., 2001) was used to mimic background synaptic inputs under a noisy *in vivo* like state. We used this point conductance model instead of a Poisson train of synaptic inputs (Williams, 2004) to ensure higher variability in the amplitude of EPSPs and IPSPs, which better reflects the background synaptic inputs with different synaptic strengths. The dynamic conductance is computed with the following differential equations (**Eq. 4-6**) (χ_1_ and χ_2_ are two independent random variables following unit normal distribution),

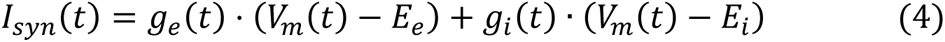

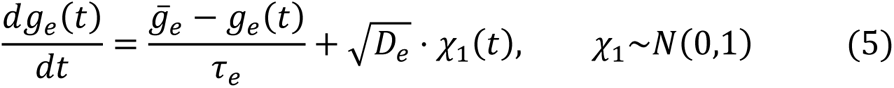

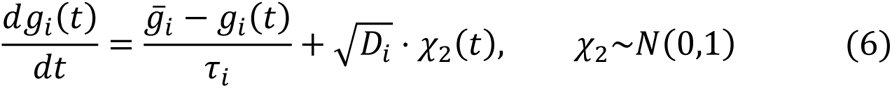

In our experiment, *τ*_e_= 2.8 ms, *τ*_i_ = 8.5 ms, E_e_ = 0 mV, E_i_ = -80mV, were chosen based on the physiological properties of α-amino-3-hydroxy-5-methyl-4-isoxazolepropionic acid (AMPA) and γ-Aminobutyric acid (GABA) channels. The mean of noisy excitatory conductance 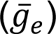 varies between 2 nS to 4 nS for different amount of spontaneous background firing rate, and 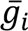 equaled one half of 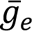. Conductance values agreed with previous literature and this range helped match the background firing rates to that observed *in vivo* (2 to 20 Hz), but avoided a hyperexcitable state. The noise diffusion coefficients of the excitatory (D_e_) and inhibitory conductance (D_i_) were scaled to match voltage fluctuations normally observed *in vivo* (∼ 5mV RMS). These terms, which contribute to the “noisy” nature of dynamic conductance, were in line with previous literature (Destexhe et al., 2001) and this ensured the fast-fluctuations in the background were preserved.

### Simultaneous two photon imaging and 3D holographic glutamate uncaging

Simultaneous two-photon imaging and 3D holographic glutamate uncaging was performed using a laser-scanning microscope (Bruker Nano, Madison, WI) fitted with dual-galvanometric mirrors, a spatial-light-modulator, custom designed pulse compression and diagnostic measurements, and two femtosecond pulsed lasers (Insight X3 and Mai-Tai, Spectra Physics). Laser beam intensities were independently controlled with a pair of electro-optical modulators (model 350-50; Conoptics, USA). A 512*512 spatial light modulator (Meadowlark Optics/Boulder Nonlinear Systems, Lafayette, CO) was coupled to the uncaging path and relayed to the pair of uncaging galvanometers. A half-wave plate was used to achieve the preferred polarization by monitoring the first order efficiency of the SLM, and a look up table optimization was performed to maximize the diffraction efficiency at 740 nm. A zero-order beam block was placed at the conjugate image plane. A custom-designed Temporal Dispersion Compensation (TDC) module in conjunction with a Michelson intensity autocorrelator was employed for pulse compression and diagnostic measurements. The TDC arm comprised of a dual Prism assembly in folded mirror configuration and the compressed pulse was then redirected by a pickoff mirror to Michelson Interferometer for pulse width measurements at the microscope sample plane. The imaging beam (810 nm for Alexa 594 and fluo-4, 920 nm for Alexa 488) and uncaging beam (740nm) were combined using a 760 nm long pass dichroic mirror. During Calcium (Ca^2+^) imaging line-scans were performed across the spine head and the adjacent branch with 8 µs to 10 µs dwell time (400 to 800 Hz line rate), and 3 mW to 5 mW laser power at the objective focal plane. Structural imaging was performed at 5 µs to 8 µs dwell time. For glutamate uncaging with 4-methoxy-5,7-dinitroindolinyl-L-glutamate trifluoroacetate (DNI-glu-TFA) (Femtonics inc., Budapest, Hungary) (Tønnesen et al., 2014), 1.5 mM to 2 mM DNI-glu-TFA was diluted in freshly prepared recording ACSF and applied to the bath through a circulating pump. At these concentrations there was no epileptiform-like activity. The uncaging dwell time was 1 ms and the laser power needed for uncaging ranged from 12 mW to 18 mW. At these powers no visible photodamage occurred. Baseline fluorescence of both channels was continuously measured to assay any damage. Ca^2+^ transients were also measured to ensure spines were still functionally active with no loss in physiological response. EPSP time-course and changes in resting membrane potential following repeated stimulation were also assayed as indicators of any photodamage. Although undesired bleed-through of maximally extinct laser power at the sample plane was <1mW, table-top hard shutters were used to avoid exposure and any off-target uncaging. The minimum time required for the SLM to update its phase mask was 3 ms which also set the limit for how fast multiple synaptic clusters spread across different arbors could be stimulated. Ca^2+^ signals were expressed as DF/F (calculated as (F - F_baseline_)/F_baseline_). Data was collected from dendrites that were at least 30 µm below the surface of the slice that were not prematurely cut off before termination.

### PSF Estimation of Holographic Uncaging Spots

We estimated the holographic uncaging resolution of our customized SLM microscope experimentally by imaging sub-resolution fluorescent beads (100 nm FluoSpheres, Molecular Probes, USA) embedded in 2 % agarose (Cole et al., 2011) when SLM was engaged. The excitation wavelength of the pulsed laser was set at 800 nm. Serial sections of the sample were performed at a step size of 100nm. Intensity profiles along horizontal and vertical to the image plane through the bead were measured. After subtracting the baseline fluorescence intensity measured from the dark background, the fluorescence intensity was fitted to a Gaussian (Eq. 7) (a, b and c are the fitting parameters), and the full width at half maximum (FWHM) (Eq. 8) was taken as a measure for the optical resolution of the system in lateral and axial dimension respectively (Figure 1A).

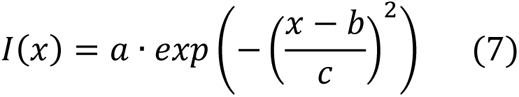

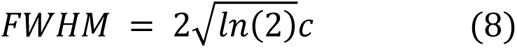

### Focal synaptic stimulation

Focal synaptic stimulation was performed with double-barrel theta pipettes (Sutter Instruments, CA, USA) filled with recording ACSF and 25uM Alexa 594. Once the neuron was patched and back-filled with the structural indicator, the theta pipette was moved to approximately 2 µm’s from the dendrite of interest under the guide of two-photon imaging. A biphasic pulse with 0.1ms duration, each phase under constant voltage mode, was used for stimulation. The stimulation intensity was gradually increased to approximately 10V to evoke unitary EPSP, verified by 0.5mV somatic EPSP amplitudes recorded via the whole-cell patch electrode. To evoke NMDAR spikes, two biphasic pulses at 100Hz were applied while gradually increasing the intensity (16 to 20V) until a nonlinear increase in EPSP amplitude was observed. To avoid the effect of inhibitory synaptic inputs, picrotoxin (0.1mM) was applied to the bath.

### Pharmacology

D-AP5 (Tocris, USA) was suspended in water as 50 mM stock solution. Tetrodotoxin (TTX, Tocris, USA) was suspended in 2 mM citric acid as 1mM stock solution and diluted to 1 µM with ACSF on the day of experiment. For focal synaptic stimulation, 0.1 mM picrotoxin (Tocris, USA) was dissolved in ACSF in a warmed water bath under constant stirring on the day of experiment. Recording ACSF containing pharmacological blocker(s) was washed in through the perfusion system. Recordings were performed 5 to 10 minutes after pharmacological blocker wash-in.

### Surgery procedures

All experimental procedures pertaining to animal surgeries and in vivo experiments were conducted in accordance with the guidelines set forth by the NIH and Purdue Institutional Animal Care and Use Committee (IACUC). C57BL/6 male and female mice were used in approximately equal numbers. Adult mice (between 8 and 24 weeks of age) were used for in vivo experimentation. Mice were kept on a 12-hour light/dark cycle in conventional housing and had unrestricted access to food and water. Adult C57BL/6 mice were deeply anesthetized with 3 to 4% isoflurane. Anesthesia was maintained with 1 to 1.5% isoflurane during surgery with oxygen flow rate ∼0.1 L/minute. A thermo pad (Kent scientific) was used during the surgery to maintain the body temperature. Carprofen and Dexamethasone (0.6mg/kg body weight) was injected subcutaneously, and lidocaine injected under the scalp after the induction of anesthesia. Eye ointment was applied, and the scalp shaved and sanitized before the incision of the scalp. 3% hydroperoxide is applied on the skull and removed by dry cotton swabs to remove the excessive tissue. For intrinsic imaging, the skull was thinned to a 4 mm diameter so that the vasculature could be visualized under the stereoscope. Vetbond (3M, Saint Paul, MN) was applied to the dried skull surrounding the thinned region. A custom-designed Titanium headplate was glued to the skull with metabond (Parkell, Brentwood, NY). Mice were injected with carprofen Intraperitoneally for up to two days after the surgery. Prior to silicon probe recording sessions, mice were headfixed and habituated on a circular running treadmill for 2 sessions (40 minutes each).

### Intrinsic optical imaging

Mice were head fixed, anesthetized with 3 to 4% isoflurane, and placed under a CCD camera (Blackfly USB3, Teledyne FLIR) equipped with a focus lens (Nikon, USA). The camera and lens were angled at 10 degrees for a perpendicular visualization of the barrel cortex. Light anesthesia was maintained with 0.5% isoflurane and the oxygen flow rate adjusted to 0.2L/ minutes to maintain relatively high respiration rates. This ensured an increase in the oxygenation of hemoglobin and the intrinsic image contrast. A thermo pad was used to maintain the body temperature during anesthesia. Mineral oil was applied along the thinned skull and a coverslip placed on the top to ensure a stable moisture-free image intensity. An image of the vasculature was taken as a reference with green LED illumination. A red LED (lumincor, spectral X) was used to illuminate the thinned scalp for intrinsic imaging. Custom-written Matlab (MathWorks, Natick, CA) and Arduino codes were used for intrinsic imaging. During each imaging trial, the whisker of interest was deflected with vibration of a hook controlled by a stepper motor, and 100 frames were recorded before and during the whisker deflection at 30Hz. The frames taken before and during whisker deflection were averaged respectively, and the difference between the two averaged images formed the intrinsic optical signal for one trial. The trial was repeated for 6 to 8 times, and the image from each trial was averaged until the contrast of the image was clear and the barrel distinguishable from the background. The image was then contrast-enhanced and smoothed before overlaying on top of the reference image of the vasculature.

### *In-vivo* silicon probe recording

64 channel silicon probes (from Masmanidis Lab in UCLA, 64D) were electroplated (Intan, USA) so that the impedance was lower than 200 kΩ prior to each recording session. After anesthetizing the animal, a burr hole (less than 1mm in diameter) was made with a dental drill on the skull above the target barrel, identified via intrinsic optical imaging. The silicon probe was inserted at an angle of 10 degrees and a speed of 10 µm/second. The mouse was head fixed on a circular running disk while the whisker was brushed with a post controlled by a stepper motor (Wantai motor, China). Treadmill rotation was captured by a rotary encoder (H5-1000-IE-D, US digital). Whisker deflection was performed in 15s intervals, and 200 to 400 trials of whisker deflections were performed across each animal. Whisker motion was recorded via a 45-degree mirror mounted under the treadmill onto a high-speed camera (DR1-D1312IE-200-G2-8, Photonfocus, Lachen, Switzerland) at 250 to 500 fps. The electrophysiological recording was performed with an Intan head stage (RHD2164, Intan Tech., USA) and digitized at 20 kHz. Each whisker deflection trial was truncated from 3s before to 3s after the whisker deflection. The spike sorting of silicon probe recording was performed with kilosort (Pachitariu et al., 2016) followed by manual clustering. For the local field potential (LFP), the raw data recorded from the silicon probe was filtered with a low pass filter at 500Hz and two notch filter at 120Hz and 60Hz respectively in Matlab (MathWorks, Natick, CA).

### Point-source stochastic Hodgkin-Huxley model

To simulate the probabilistic nature of ion channels, a 13-state discrete Markovian kinetic scheme (8 states for Na^+^ channels, 5 states for K^+^ channels) (Chow and White, 1996) was used and incorporated into the Hodgkin-Huxley model (stochastic HH model). This model included 3 identical activation gates (m) and 1 inactivation gate (h) for Na^+^ channels, and 4 identical activation gates (n) for K^+^ channels. The opening and closing of each binary gate were assumed to be independent and the rate of transitioning between the opening and closing state equals the opening (α) and closing rate (β) in the classic Hodgkin-Huxley model. The transition of ion channels between states can be regarded as a large numbers of independent random binary events, therefore the number of channels transitioning from state i to state j (Δn_ij_) within the time duration of Δt can be estimated with random variables that follow the binomial distribution *Δn_ij_*~*binom*([*n_i_*]),*p* = *A_ij_dt*). Here, [n_i_] denotes the number of channels currently in state i, and *A_ij_* denotes the rate of a single ion channel transiting from state i to j (**Figure S6A**, noted on the arrows that denotes transition between states). An ion channel is assumed to be conductive only in the state where all the activation gates and the inactivation gate (if applicable) were opened (n_4_ state for potassium channels, m_3_h_1_ state for sodium channels). The total Na^+^ and K^+^ ionic conductance was estimated with the multiplication of the single-channel conductance and the number of fully activated channels *g_K_* = *γ_K_* * [*n*_4_] and *g_Na_* = *γ_Na_* * [*n*_3_*h*_1_], in which γ_K_ and γ_Na_ denotes the conductance of a single fully conductive ion channel. The following ordinary differential equations (ODEs) (**Eq. 9-12**) were used in the simulations:

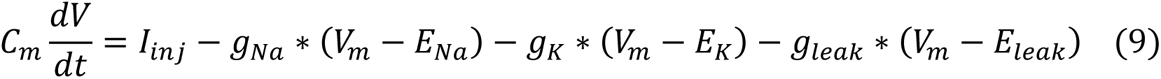

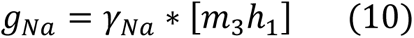

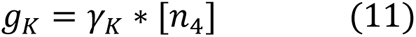

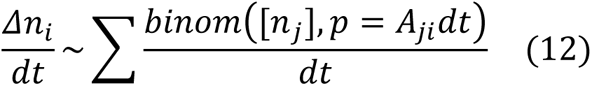

The differential equations were solved using the forward Euler method with a time interval (dt) 0.01ms. All the ion channels were at the lowest state (n_0_ for K channels and m_0_h_0_ for Na channels) at initiation, which resulted in less than 5 ms of initialization artifact, a fraction of the 250 ms period per simulation. The total numbers of Na^+^ and K^+^ channels were 12000 and 3600 respectively, other parameters can be found in **Table. S1**. The simulation was performed using custom-written python code.

## QUANTIFICATION AND STATISTICAL METHODS

The spike sorting of silicon probe recording is performed with Kilosort (Pachitariu et al., 2016), followed by manual clustering. The whisker tracing was performed with Deeplabcut (Mathis et al., 2018). The structural images were processed with ImageJ, Kalman filtered, and Fourier-based filtered when necessary. The Ca^2+^ line-scans signal was preprocessed with a Fourier-based narrowband filter to remove line noise when necessary, and the signal fitted with a double exponential curve. No smoothing was performed. Data analysis and statistical tests were performed using custom-written MATLAB (MathWorks, USA) and python code. Statistical analysis details can be found in the text and figure legends. Unless stated otherwise, the error bars represent mean +/- standard error of mean (SEM).

