## Supplemental Information for "Probing Action Potential Generation and Timing under Multiplexed Basal Dendritic Computations Using Two-photon 3D Holographic Uncaging"

**TITLE:**

This document contains Supplementary Information comprising **Figures S1 to Figure S8, Table S1, and Supplemental Notes 1**

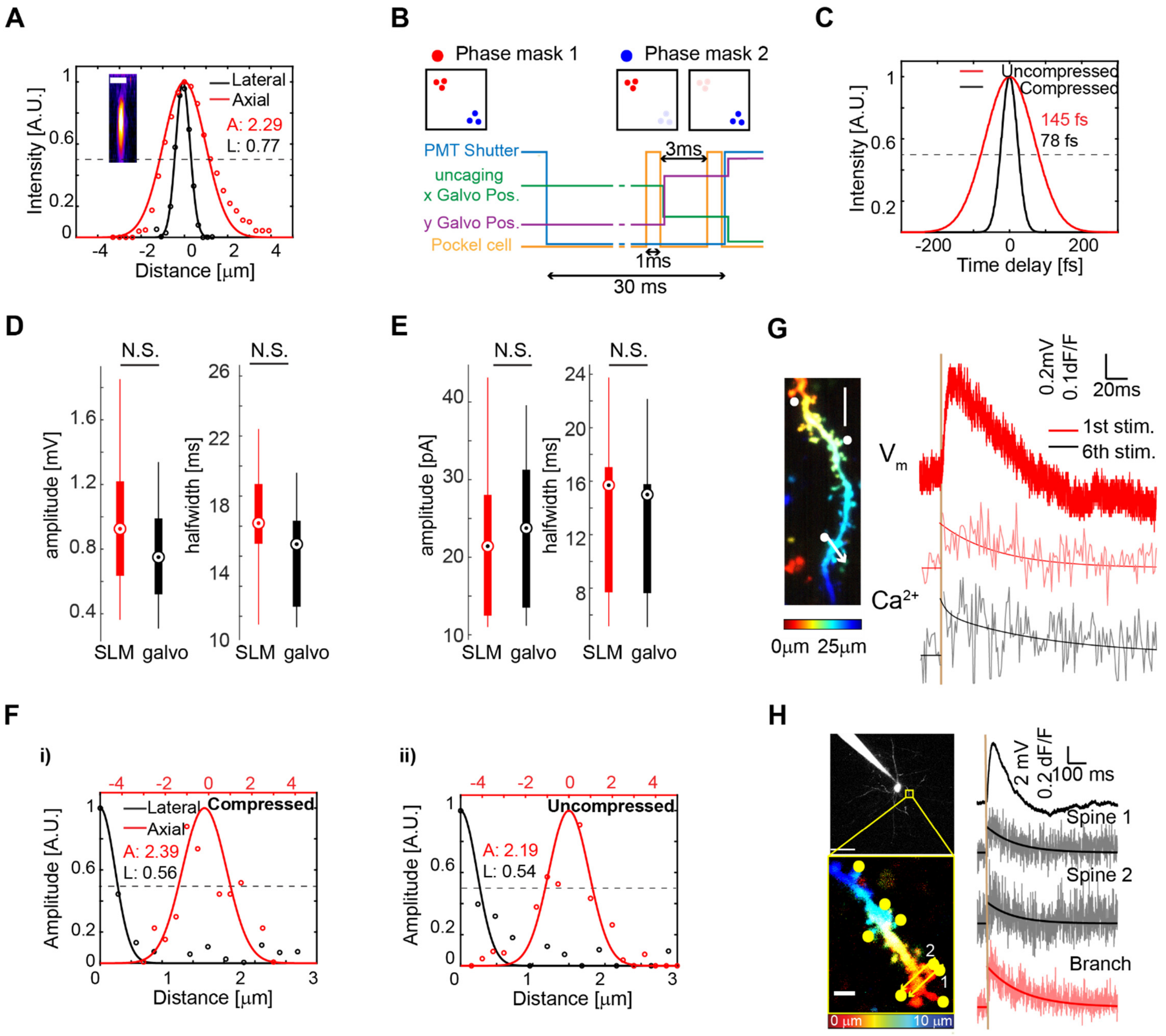


**Figure S1. Characterization of SLM-based uncaging. Related to Figure 1. A)** Point spread function under galvanometric uncaging measured with 100nm beads (inset, scale bar 2 μm), fitted with a Gaussian function. **B)** Uncaging timing diagram with simultaneous two-photon imaging. To update the SLM phase mask, a 3ms minimum interval is given for hardware responding time. **C)** Pulse width of the galvanometric uncaging beam. **D-E)** Uncaging-evoked response across 2 spines measured in terms of EPSP's **(D)** and EPSC's **(E)** were comparable with both galvanometric and SLM-based uncaging**.** **(D)** Comparison of the amplitude (left, $p = 0.07$) and half width (right, $p = 0.60$) of EPSP's evoked with galvanometric and SLM-based uncaging shows no significant difference (Paired sample student t-test, N = 17 pairs of spines). **(E)** Comparison of the amplitude (left, $p = 0.43$) and half width (right, $p = 0.71$) of EPSC's evoked with galvanometric and SLM-based uncaging shows no significant difference (Paired sample student t-test, N = 11 pairs of spines). **F) i)** Physiological PSF of galvanometric uncaging beam with pulse width compression. **ii)** Physiological PSF of galvanometric uncaging beam without pulse compression. The physiological PSF of the compressed and uncompressed beam is comparable. **G)** No photodamage or functional change was observed after repetitive uncaging. Red: Membrane voltage and initial spine head $Ca^{2+}$ transient after SLM-based uncaging across 3 distributed spines (only one used for calcium transients). Black: spine head $Ca^{2+}$ assayed after the $6^{th}$ repetitive SLM uncaging trial. Scale bar 5 µm. **H)** Branch $Ca^{2+}$ transients evoked by SLM uncaging across 8 clustered spines. Scale bar 50µm (top) and 2µm (bottom).

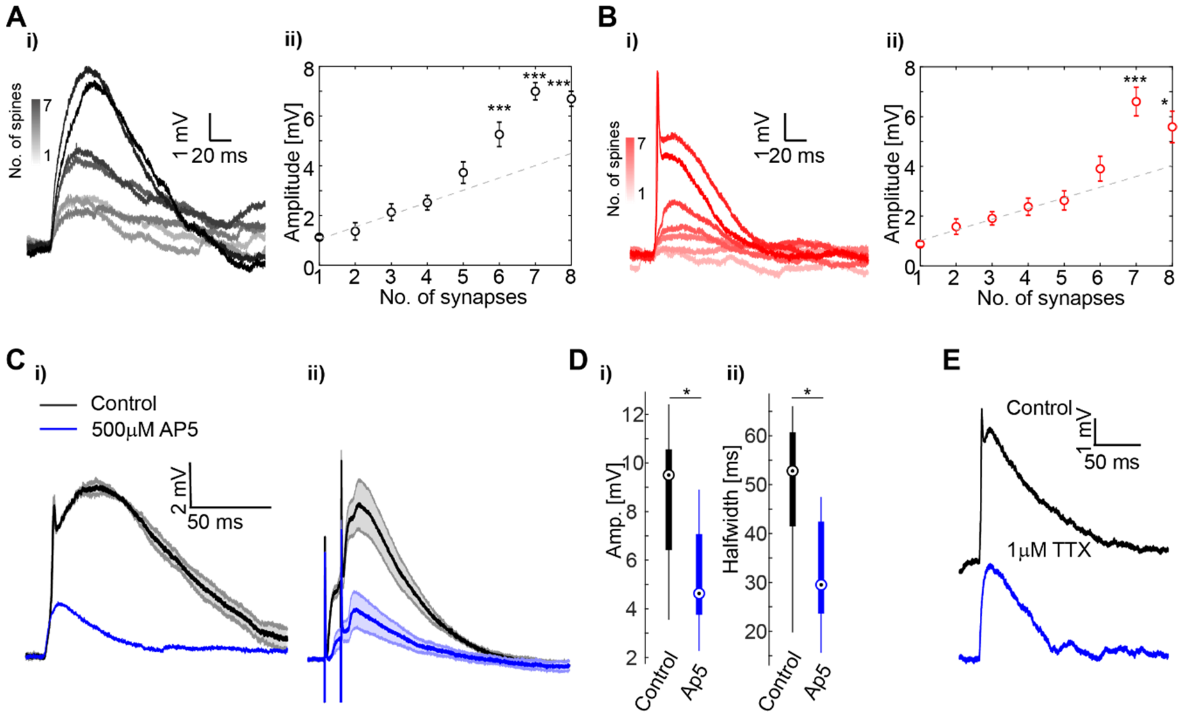


**Figure S2. Characterizing the dendritic NMDA and Na$^+$ spike response. Related to Figure 2. A)** Representative weak cluster response revealing a NMDA-only plateau response **(i)** and the resultant input-output transformations **(ii).** Note the supralinear summation when 6 to 8 synapses are co-activated (Wilcoxon signed-rank test, N = 11 weak clusters, one asterisk: $p < 0.05$, two: $p < 0.01$, three: $p < 0.005$). **B)** Similar to A, but for strong clusters with a Na$^+$ driven plateau potential (Wilcoxon signed-rank test, N = 18 strong clusters, one asterisk: $p < 0.05$, two: $p < 0.01$, three: $p < 0.005$). **C)** Basal dendritic cluster uncaging-evoked **(i)** and focal theta stimulation-induced **(ii)** plateau potentials are abolished by bath application of AP5. **D)** The amplitude and half width of the plateau potentials are clearly decreased by AP5 application. (Wilcoxon signed-rank test, N = 7, $p < 0.05$). **E)** Strong Cluster uncaging-evoked response under control (black) and 1μM tetrodotoxin (blue) application.

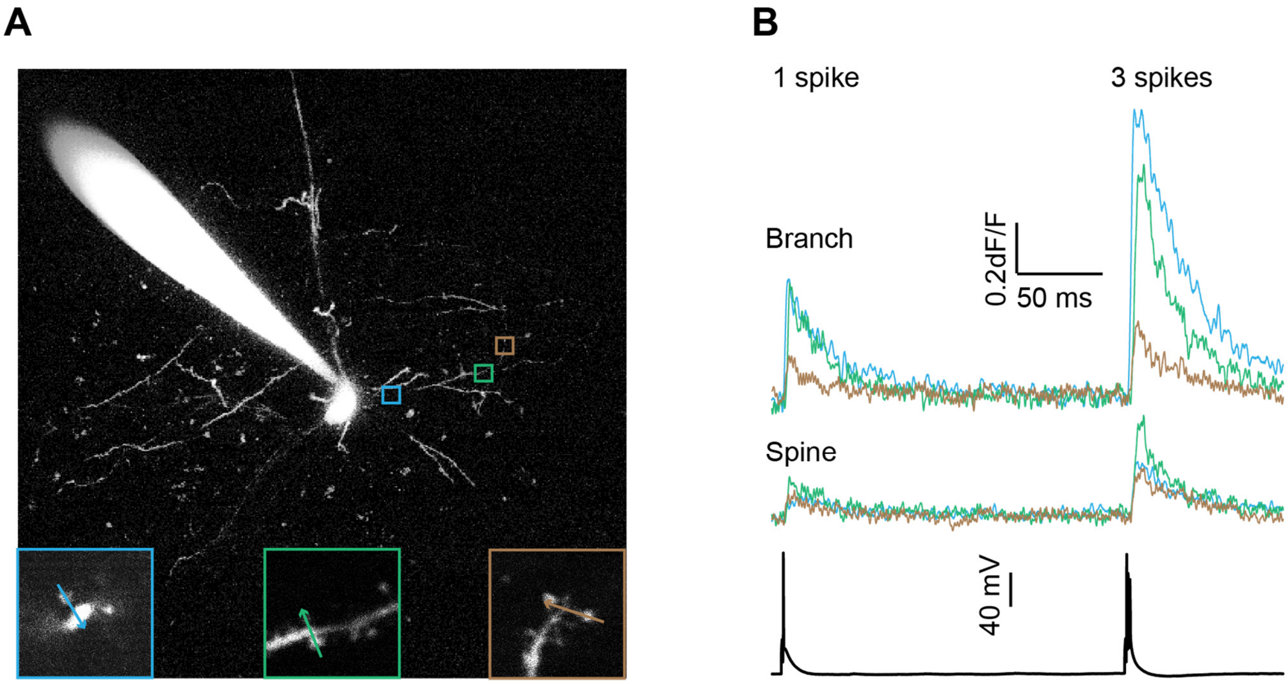


**Figure S3. Back propagating action potentials (bAPs) invade further down into distal basal dendrites and spines in a frequency dependent manner. Related to Figure 2. A)** An example L5 PN with 3 spines along the same basal dendritic branch located at different distances from the cell body, scale bars, 20 μm. Insets: zoomed-in view of the spines of interest, the arrow indicates the line scan, scale bars, 2 μm. The distance from the cell body (from left to right) is 25μm, 72μm, 90μm respectively. **B)** $Ca^{2+}$ response on the dendritic shaft (top) and spine head (middle) evoked by single action potential and a triplet burst. Note the $Ca^{2+}$ signal is stronger in both the spine head and dendritic shaft when evoked by triplet compared to a single AP.

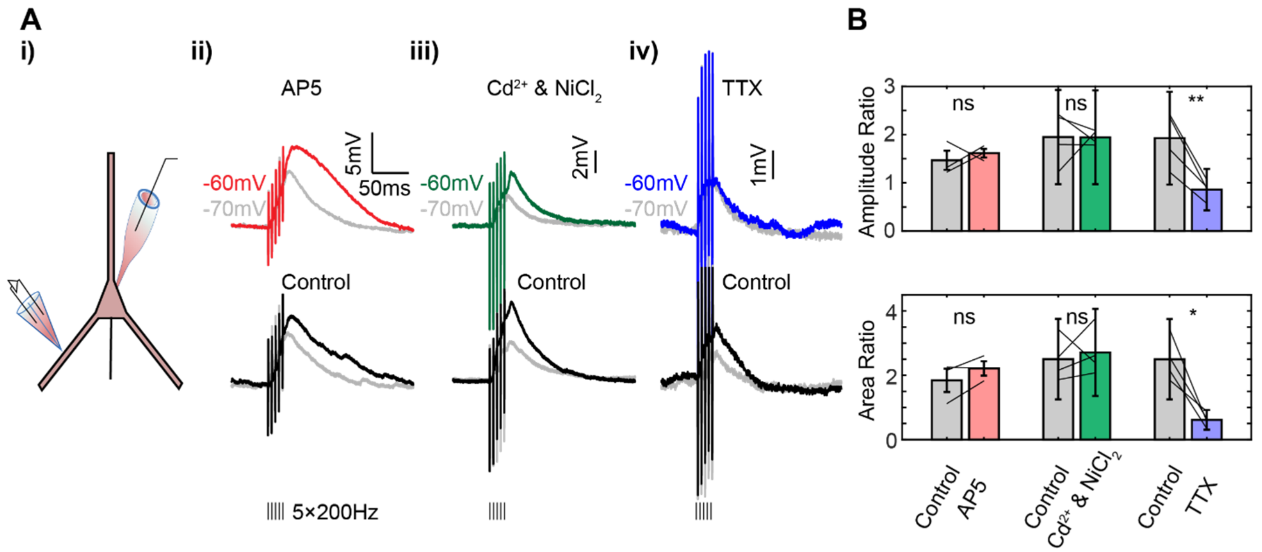


**Figure S4. Voltage-dependent amplification of synaptic integration in the subthreshold range is mediated by $Na^+$ channels. Related to Figure 3. A) i**) Experimental configuration wherein a theta pipette is used to evoke EPSPs in the basal dendrite of L5 PN. ii) Modulating the RMP amplifies the somatic response to a train of 5 theta-stimulations at 200Hz, under control (black) and application of 100µM AP5 (red). iii) Similar to ii), but under control (black) and bath application of 100µM $CdCl_2$ and 100µM $NiCl_2$ (green). iv) Bath application of 50nM TTX abolishes the voltage-dependent amplification. **B)** The amplification ratio (top) and integral (area-under-curve, bottom) upon modulation of the RMP (-60mV/-70mV) under control and different drug applications (Paired sample t-test, AP5: N = 3 neurons, $Cd^{2+}$ and $Ni^{2+}$: N = 4 neurons, TTX: N = 4 neurons, $p < 0.01$ for amplitude, $p < 0.05$ for area-under-curve).

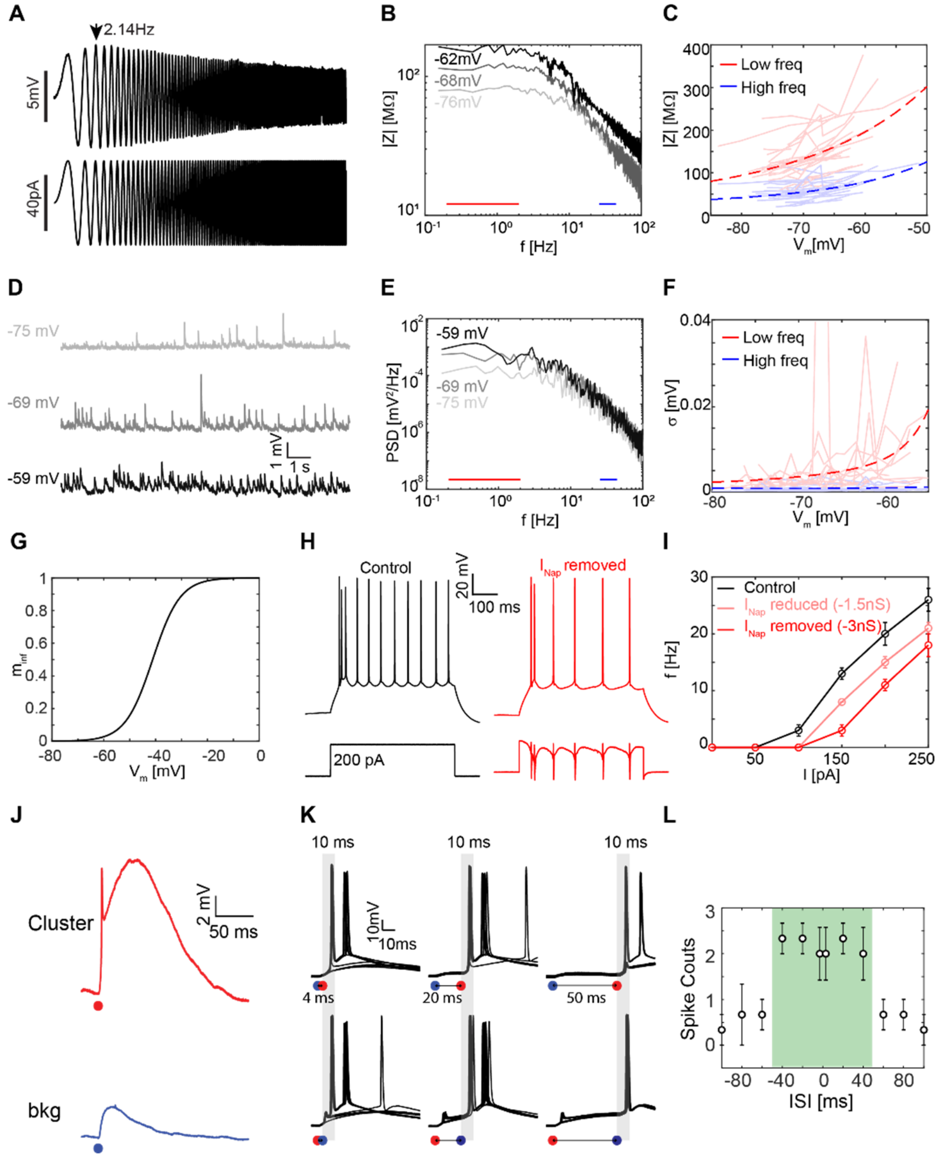


**Figure S5. Increased impedance with subthreshold depolarization amplifies weaker inputs and enhances gain. Related to Figure 3. A)** Somatic injection of Zap current (bottom) was used to measure the impedance of L5PN. Arrow indicates the resonance frequency of L5 PN cell body. **B)** Impedance spectra of a L5 PN plotted at different resting membrane voltage (RMPs). The red and blue line marks the frequency bands used in **(C)**. **C)** Impedance across low (red, 0.5 to 2 Hz) and high (blue, 15 to 30 Hz) frequency bands as a function of RMP, fitted with exponential function (N = 25 neurons). **D)** An example of spontaneous subthreshold $V_M$ fluctuations recorded from a L5 PN at different resting membrane voltages. **E)** Power spectral density (PSD) of $V_M$ at different RMPs. The red and blue line indicates the low and high frequency bands used in **(F)**. **F)** $V_M$ power at low (red, 0.5 to 2 Hz) and high (blue, 15 to 30 Hz) frequency bands as a function of resting $V_M$, fitted with exponential function (N = 13 neurons). **G)** The sigmoidal steady-state opening gate parameter ($m_{inf}$) of $I_{NaP}$ channels used in the dynamic clamp experiments to remove $I_{NaP}$ conductance. **H)** Removing $I_{NaP}$ ($G_{Nap}$ = -3 nS) conductance with the dynamic clamp decreases the excitability of the L5 PN. **I)** f-I curve of L5 PN with subtraction of 1.5nS (pink) and 3nS (red) $I_{Nap}$ conductance with dynamic clamp (N = 3 neurons). **J)** Uncaging response of a cluster A (red) and 5 background synapses (blue) on a L5PN. **K)** Sequential uncaging of the cluster A and the distributed background synapses (marked with red and blue circle) with different ISI shows a time window of +/- 50 ms where the cluster cooperates with the synaptic background and evokes doublets (10ms window after the uncaging event is marked with gray shadow). With longer temporal separation between uncaging cluster A and the background depolarization, uncaging-evoked action potential probability goes down (not shown). **L)** Number of spikes evoked as a function of ISI between two group uncaging events. The green shadow marks the +/- 50 ms integration window where the somatic AP output gain is increase (N = 3 neurons).

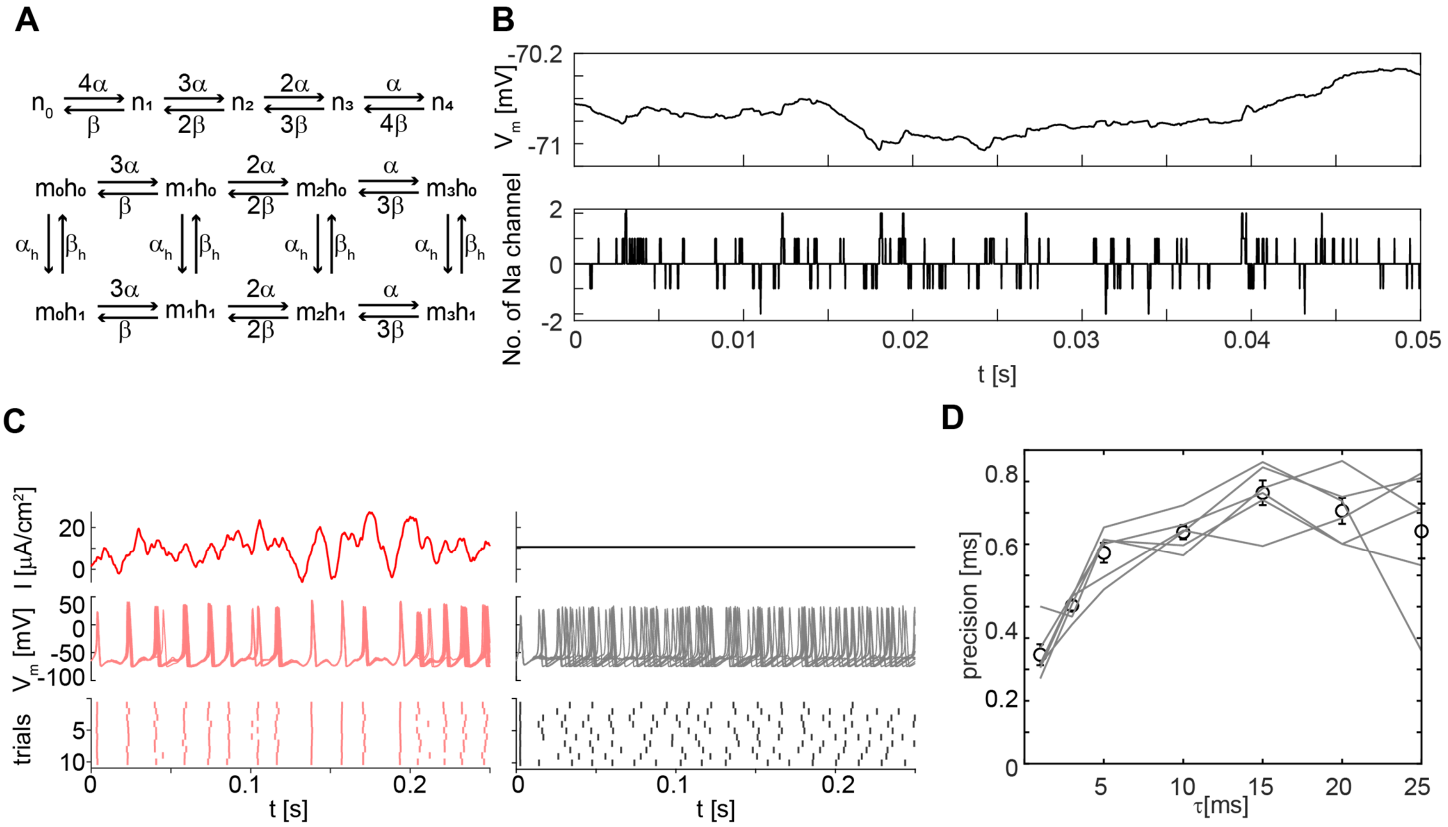


**Figure S6. Stochastic ion channel gating reveals an increase in temporal precision under dynamic current injection. Related to Figure 4. A)** The stochastic gating Hodgkin-Huxley model. The 13-state kinetic HH model used for simulating stochastic ion channel gating. **B)** Subthreshold $V_M$ fluctuation as a result of stochastic gating (top), and the corresponding number of fully-opened $Na^+$ channels (bottom). The opening and closing of ion channels are stochastic in nature, which creates spontaneous subthreshold voltage noise. **C)** Somatic injection of dynamic current (red) and DC current (black) to the stochastic HH model in **(A)**. The dynamic current evokes somatic spike with higher precision. **D)** Spike temporal precision as a function of the time constant of the injected current, N = 6 independent simulations.

**Table S1: Parameters in the stochastic HH model** (Adapted from (Schneidman, Freedman, & Segev, 1998).

| parameter type | value |
|---|---|
| dt | 0.01 ms |
| $C_m$ | 1 µf/cm$^2$ |
| $E_{Na}$ | 50 mV |
| $E_K$ | -77 mV |
| $E_{leak}$ | -54.4 mV |
| $\gamma_K$ | 20 pS |
| $\gamma_{Na}$ | 20 pS |
| Density_Na | 18 channel/µm$^2$ |
| Density_K | 60 channel/µm$^2$ |
| $\bar{g}_{leak}$ | 0.3 mS/cm$^2$ |
| area | 200 µm$^2$ |

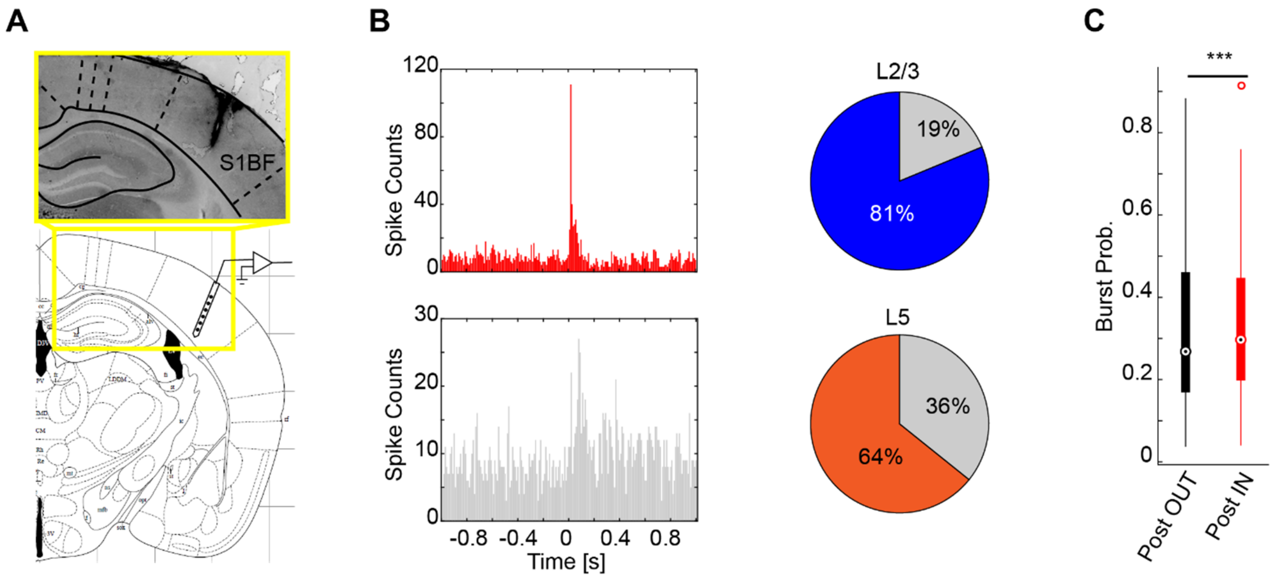


**Figure S7. *In vivo* silicon probe recordings in S1. Related to Figure 5. A)** Probe insertion track in the Barrel cortex. **B)** Left: Example peri-stimulus time histogram (PSTH) for a representative "stimulus-locked" single unit (red) and a single unit that does not show a strong time-locked response to whisker stimulation (gray). Right: Pie charts showing the percentage of "stimulus-locked" single units in L2/3 (blue) and L5 (orange). Majority of single units in L2/3 and L5 are time-locked to the whisker stimulation (N = 32 single units in L2/3 and 137 single units in L5 from 6 animals). **C)** 30% of L5 single unit activities are in the form of high frequency bursts (50Hz or higher). (Wilcoxon signed-rank test, N = 137 L5 single units from 6 animals, $p < 0.001$).

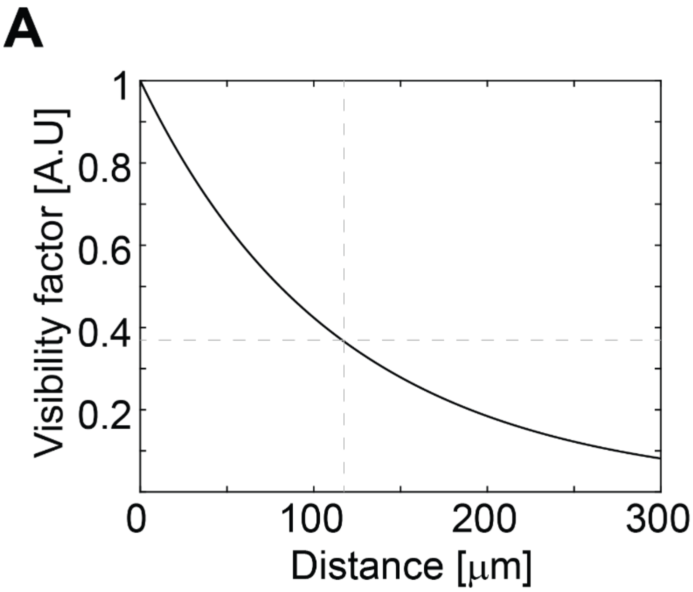


**Figure S8. Conductance visibility in dendrites. Related to Figure 5. A)** Steady state conductance visibility factor in a passive basal dendrite (See **Supplemental Note 1** for details).

### SUPPLEMENTAL NOTE 1

#### Analytical estimation of conductance visibility in the dendrites

Following the method outlined by Koch and colleagues (Koch, Douglas, & Wehmeier, 1990), we estimated the visibility factor of the somatic high conductance state in the basal dendrites. We assumed the dendrites are passive, in which case the visibility factor can then be expressed with:

$$\Gamma = \frac{exp\left(-\frac{2x}{\lambda_{ds}}\right)}{1 + \left(1 - exp\left(-\frac{x}{\lambda_{ds}} - \frac{x}{\lambda_{sd}}\right)\right) K_{ss} g_s}$$

Here, $\lambda_{ds}$ is the forward propagation (dendrite to soma) space constant, $\lambda_{sd}$ is the back-propagation space constant, $K_{ss}$ is the somatic input impedance, $g_s$ is the somatic conductance, and x is the distance of the dendrite segment to the soma. When the somatic conductance introduced by the dynamic clamp is relatively lower than the somatic membrane conductance ( $K_{ss} g_s \ll 1$ ), the visibility factor approximately follows the exponential decay, with the space constant about half of the forward propagation space constant ($\frac{\lambda_{ds}}{2}$). Under this condition, the dendritic space constant ($\lambda_{ds}$) is the major determining factor of conductance visibility, suggesting the conductance visibility increase with higher dendritic input impedance. Using results from published experimental findings (Nevian, Larkum, Polsky, & Schiller, 2007), the steady state forward propagation space constant was assumed to be 250μm, and the steady state backward space constant was assumed to be 609μm. The somatic resistance used was within our experimental range of ~70MOhm, and the average somatic dynamic conductance was 2nS. With these values the half-decay point of conductance in the dendrite was around 80μm, while the space constant for conductance visibility was ~ 120μm.
